# Ab-trapping - a peripheral staining artifact in antibody-based microscopy and genomics

**DOI:** 10.1101/2025.04.09.648027

**Authors:** Konrad Chudzik, Yuko Sato, Xingchi Yan, Simon Ullrich, Watanya Trakarnphornsombat, Lothar Schermelleh, Geoffrey Fudenberg, Hiroshi Kimura, Michael I. Robson, Irina Solovei

## Abstract

Antibodies (Ab) are essential for detecting specific epitopes in microscopy and genomics, but can produce artifacts leading to erroneous interpretations. Here, we characterize a novel artifact, Ab-trapping, in which antibodies bind at the periphery of a cellular structure and do not diffuse further into its interior. This causes anomalous peripheral staining for multiple critical targets, including endogenous or ectopically expressed nuclear proteins like nucleolar proteins, histone variants and their modifications like H3K9me2. Ab-trapping can affect any assay relying on Ab diffusion, including immunofluorescence microscopy and recent genomics approaches like CUT&Tag. Critically, computational modeling and experimental validation reveal that Ab-trapping is caused by high epitope abundance, high Ab affinity, and low diffusion rates. Consequently, its effects can be mitigated by using alternative Abs and optimizing incubation conditions. Ab-trapping is therefore a considerable artifact that should be considered when designing experiments and interpreting results.

## MAIN

Antibodies (Abs) are key tools in microscopy and genomics for determining how epitopes are distributed throughout the cell and across the genome. However, Abs can cause artifacts due to multiple factors, including cross-reactivity to undesirable targets, epitope masking, and fixation-induced changes in protein distribution^1–6^. Ab artifacts can thus potentially bias both microscopy methods and genomics approaches, all of which rely on applying Abs to intact cells to detect chromatin-associated proteins, including CUT&RUN/ChIC, CUT&Tag, pA-DamID and MAbID^7–11^. The above genomics approaches also underlie emerging state-of-the-art technologies that map epitope distributions in whole single cells^9,11–13^ or in space within intact tissues^14^. Being aware of potential Ab artifacts and methods to mitigate them is thus critical to prevent false interpretations from classical and more recent molecular methodologies in numerous fields in biological research.

Here, we describe “Ab-trapping”, a prevalent yet previously uncharacterized artifact caused by the trapping of Abs on the edge of cellular structures, resulting in artifactual peripheral signal enrichment. We demonstrate that Ab-trapping impacts assays like immunofluorescence microscopy and CUT&Tag sequencing. We observed this effect for multiple Abs against several critical targets, including transcription factors and histone modifications in the nucleus and proteins in the nucleolus. Using computational modeling and experimental validation, we identified three key parameters that synergistically influence Ab-trapping - epitope abundance, Ab affinity, and Ab diffusion dynamics. Low epitope concentration, low Ab affinity, and high diffusion coefficients facilitate antibody penetration, resulting in interior staining. Conversely, the opposite parameters display peripheral staining caused by Ab-trapping. Together, our data demonstrate that Ab-trapping is a generalisable phenomenon that is relevant to many Ab-based assays and must be considered to avoid incorrect conclusions.

## RESULTS

### Anti-H3K9me2 Abs produce conflicting signals in standard immunofluorescence

The spatial segregation of active euchromatin from inactive heterochromatin is required for the proper function of the cell nucleus^15–17^. Accurately mapping the distribution of heterochromatin is therefore essential for interpreting nuclear organization. We thus set out to systematically profile the spatial positioning of a crucial, often-studied component of heterochromatin, histone H3 dimethylated on lysine 9 (H3K9me2), using standard immunofluorescence microscopy and three different anti-H3K9me2 Abs. These included two commercial Abs, Active Motif AM39239 and Abcam ab1220, and the CMA317 Ab generated in the Kimura laboratory^18^.

We first co-stained mouse myoblasts and human HeLa cells with rabbit AM39239 mixed with either mouse CMA317 or mouse ab1220. Both mouse Abs produced staining that was consistently distributed throughout nucleoplasm, including the nuclear periphery, the periphery of nucleoli, and other loci (Fig.1a). However, AM39239 was strikingly confined to the nuclear periphery with only a residual gradient of staining towards the nuclear interior. Quantifying the radial signal distribution confirmed these contradictory staining results, validating that the signal of AM39239 is more peripherally enriched than the signal from the other two anti-H3K9me2 Abs (Fig.1b). Moreover, these distinct staining patterns of the three anti-H3K9me2 Abs proved highly consistent. AM39239 staining was persistently more restricted to the periphery than CMA317 or ab1220, both when each Ab was used separately (Extended Data Fig. 1a) and when applied together across multiple cultured cell-types (Extended Data Fig. 1b). Moreover, we find similar differences at different stages of the cell cycle. While CMA317 or ab1220 stained through condensed mitotic chromosomes during anaphase, AM39239’s signal was almost entirely restricted to their periphery (Fig.1c).

**Figure. 1.**
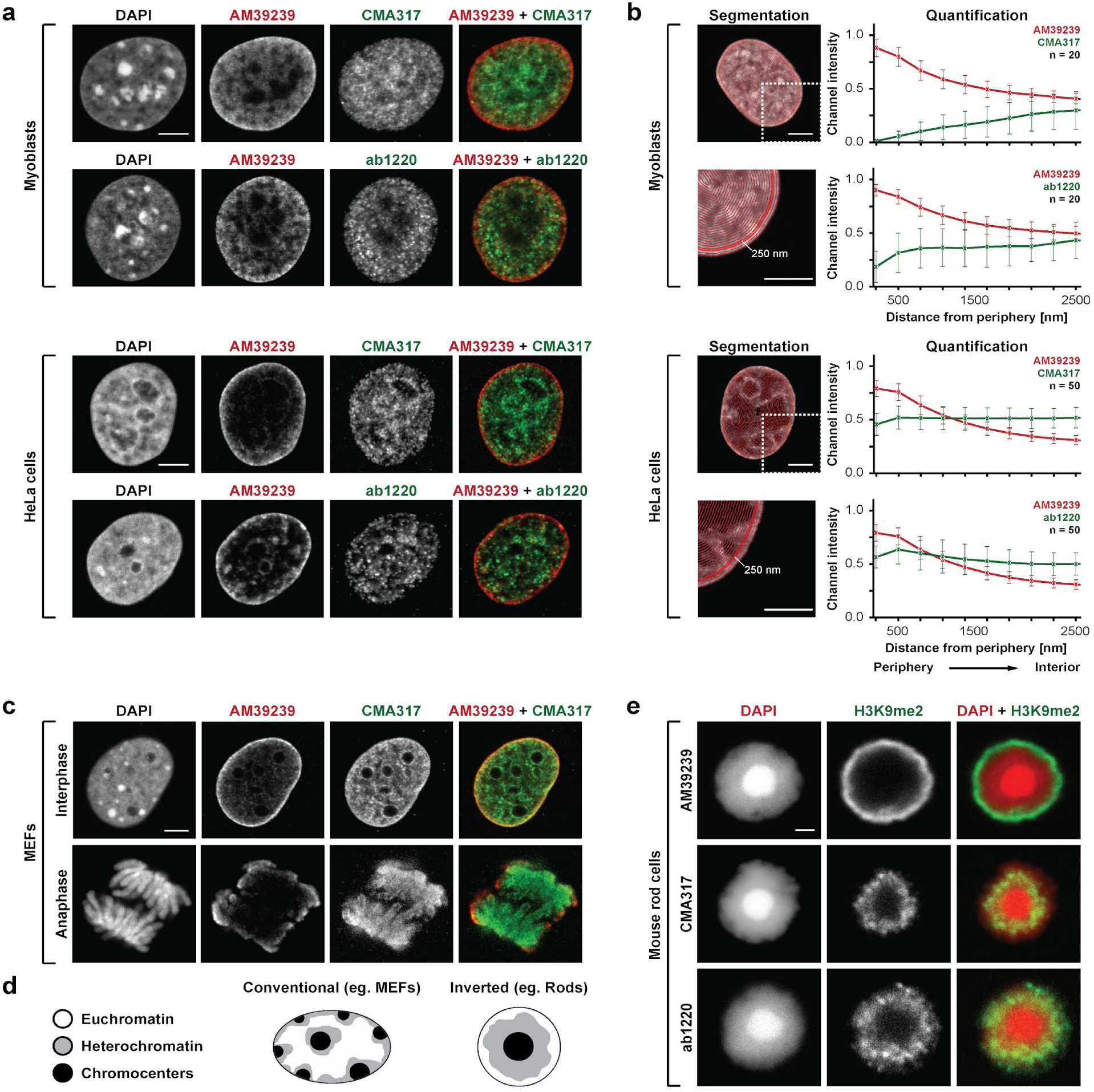
Anti-H3K9me2 Abs display contradicting immunofluorescence staining patterns. **a**, Immunofluorescence staining with pairs of distinct anti-H3K9me2 Abs in cultured mouse (myoblasts) and human (HeLa) cell nuclei. **b**, Quantification of immunofluorescence staining shown in a. Graphs show distribution of Ab signal intensities in concentric shells of AM39239 (red curves) and ab1220 or CMA317 (green curves). Nuclear segmentation into shells is depicted on the left with red contour lines. For details of nuclear segmentation and intensity normalization see Methods. n, number of analyzed nuclei. **c**, Immunofluorescence staining with pairs of anti-H3K9me2 Abs in interphase and mitotic cells in cultured mouse embryonic fibroblasts. **d**, Schematics of chromatin organization in conventional and inverted nuclei. **e**, Immunofluorescence staining with single anti-H3K9me2 Abs in isolated inverted rod nuclei. All images are single optical sections. Scale bars: a-c, 5 µm, e, 2 µm.

These contradictory Ab behaviors are also mirrored in isolated rod cell photoreceptors, a unique cell-type with a radically inverted nuclear organization where heterochromatin occupies the nuclear interior (Fig.1d) ^16,19^. Here, CMA317 and ab1220 stained the layer of heterochromatin surrounding the central chromocenter in the nuclear interior (Fig.1e). By contrast, AM39239 specifically marked the nuclear periphery, where no H3K9me2 staining is expected. Critically, these contradictory results match conflicting H3K9me2 stainings previously reported for these same Abs in rods, with CMA317 and ab1220 Abs marking the nuclear interior^20^ and AM39239 the nuclear periphery^21^.

Thus, different Abs reproducibly report highly distinct distributions of H3K9me2 in multiple contexts from conventional and inverted nuclei to mitotic chromosomes. Moreover, this contradiction is not due to interference between the three Abs, as each behaves the same whether applied individually or in combination (Fig.1a,c and Extended Data Fig. 1).

### Peripherally-restricted H3K9me2 staining is a technical artifact

We next sought to determine which staining pattern of the three anti-H3K9me2 Abs is artifactual and what causes them to behave differently. We noted three aspects of AM39239’s staining that were inconsistent with previous observations. First, the restriction of AM39239’s staining to a narrow rim of peripheral chromatin in conventional nuclei was unexpected given the reported enrichment of H3K9me2 at nucleolar-associated domains (NADs) in the nuclear interior^22^. Second, the observation of H3K9me2 enrichment in euchromatic periphery of inverted rod nuclei contradicted its established role in transcriptional repression^15^. Third, nearly exclusive peripheral H3K9me2 staining in anaphase chromosomes seemed unlikely considering that euchromatin and heterochromatin are not spatially segregated in mitotic chromosomes^23^. Combined, we reasoned that AM39239’s tendency toward peripheral accumulation is an artifact while CMA317 and ab1220 Ab staining report H3K9me2’s true distribution.

To test this, we first hypothesised that AM39239 displays unspecific binding to off-target epitopes, thereby causing incorrect staining. We thus performed immunostaining of mouse embryonic fibroblasts (MEFs) that are deficient in five H3K9 methyltransferases (5KO-MEFs) and so completely lack H3K9me2^24^. Both mouse Abs CMA317 and ab1220 displayed no signal, demonstrating that they are highly specific to H3K9me2 (Extended Data Fig. 2a). In contrast, rabbit AM39239 exhibited a residual non-specific signal throughout nuclear chromatin, corresponding with that described in the Antibody Validation Database (<u>AVD</u>). However, since AM39239’s signal enrichment at the nuclear rim was also eliminated, this indicates that the peripheral signal is at least dependent on the presence of the H3K9me2 epitope.

This indicated that the distinct staining patterns of the three anti-H3K9me2 Abs are not due to differences in epitope specificity. Instead, the differences in staining between AM39239 and the two other Abs suggests they have differing abilities to penetrate the nucleus. To examine this, we attempted to physically “break” the nuclear surface to improve the accessibility of Abs to chromatin. We generated a suspension of briefly fixed retinal cells, embedded them in a cryoblock, and prepared 14 µm thick cryosections for staining with AM39239. Cell nuclei lying entirely within the sections maintained their integrity and exhibited characteristic AM39239 nuclear rim staining (Extended Data Fig. 2b). In contrast, nuclei that were physically bisected by the sectioning, as visible by disruptions in the DAPI channel, showed staining of internal chromatin that was now exposed. This indicates that the AM39239 Ab can successfully stain heterochromatin throughout the nucleus but only when it has direct access to the nuclear interior. This finding raised the question of what could prevent the access of AM39239 to epitopes inside the nucleoplasm.

We thus investigated if the peripheral staining by AM39239 is linked to differences in its binding affinity to H3K9me2 compared to ab1220 and CMA317. We directly compared the affinities of the two monoclonal mouse Abs, CMA317 and ab1220, using ELISA. This revealed that ab1220 exhibits approximately 27-fold lower affinity than CMA317 (Extended Data Fig. 3a). Based on the previously determined binding coefficient of CMA317 (K_D_ = 1.5 × 10^-8^ M)^18^, we therefore calculate a K_D_ of ∼4 × 10^-7^ M for ab1220. We then took a stepwise approach to infer the binding affinity of the rabbit Ab AM39239 which, as a polyclonal Ab, contains a mixture of distinct IgG molecules. First, we measured AM39239’s effective concentration of H3K9me2 binding molecules. Although the total IgG concentration in AM39239 antisera was ∼5 mg/ml, only ∼0.1 mg/ml (∼4%) effectively bound H3K9me2 (Extended Data Fig. 3b-d and see Methods). Using this effective IgG concentration, we performed a competitive binding assay by applying a fixed amount of AM39239 to H3K9me2-labeled beads in the presence of increasing concentrations of CMA317. AM39239 binding was reduced by approximately 50% when competing with a ∼6-fold excess of CMA317 (Extended Data Fig. 3e). Critically, a fraction of AM39239 IgG molecules remained bound even in the presence of a ∼100-fold excess of CMA317 (Extended Data Fig. 3f). These results indicate that AM39239 contains IgGs with an average affinity that is ∼6-fold higher than CMA317, but likely spanning a broad range, including fractions with an even greater affinity.

Collectively, this suggests that the persistent peripheral staining with AM39239 is due to insufficient Ab penetration into the nucleus, which could be related to its higher binding affinity to the H3K9me2 epitope.

### Computer simulations identify parameters affecting Ab penetration

We next dissected how peripheral staining artifacts can arise using a minimal computational model that captures the essential dynamics of Ab staining. We modeled staining of the nucleus over time as a reaction-diffusion process, similar to models used to optimize tissue-scale staining^25–26^. Specifically, we approximated the nucleus as a 10 µm diameter sphere with an antigen (Ag) uniformly distributed throughout its volume. We assumed that an Ab moves according to a specific diffusion coefficient (D_A_). We also assumed Ab-Ag interactions are governed by their association (k_on_) and dissociation (k_off_) rates (Fig.2a). The baseline values for these parameters were adapted from published datasets. These included (i) the known affinity of the CMA317 Ab, (ii) the estimated H3K9me2 epitope concentration per single mammalian nucleus, (iii) and the diffusion coefficient for IgG molecules in aqueous solution (Extended Data Fig. 4a and see Methods)^18,27–29^. In these conditions, our simulations show full penetration of an Ab throughout the nuclear interior after 30 min incubation (Extended Data Fig. 4b,c).

**Figure 2.**
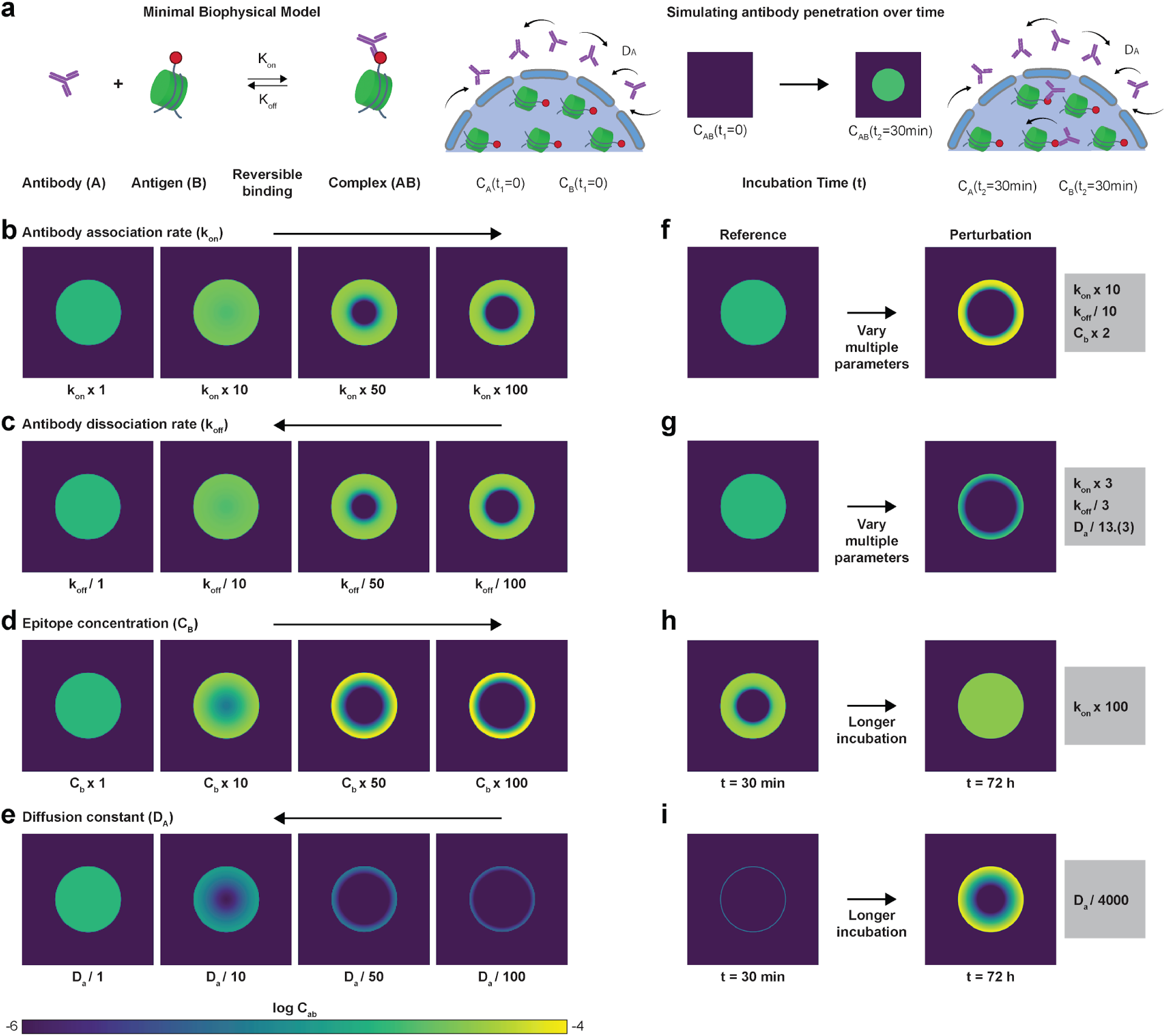
Computational modeling recapitulates peripherally restricted Ab staining. **a**, Minimal biophysical model. Left: antibody and antigen bind and unbind reversibly with rate k_on_ and k_off_. Right: at time zero, Abs are primarily located outside the nucleus, and they diffuse into the nucleus with a diffusion constant D_A_. The concentration field C_AB_ reflects this simulated distribution of bound Ab and is illustrated at time t = 0 and after a 30-minute incubation. **b-e**, C_AB_ at time t = 30 min in simulations where parameters for k_on_ (b), k_off_ (c), C_B_, (d), and D_A_ (e) are individually altered to produce Ab-trapping (See Extended Data Fig. 4). **f-g**, Trapping at time t = 30 min can also be realized by smaller alterations that affect combinations of parameters, including k_on_, k_off_ and C_B_ (f) or k_on_, k_off_ and D_A_ (g). **h**, A 100-fold increase of k_on_ leads to strong trapping at 30 min, which can be abolished with a prolonged incubation time (t) = 72 h. **i,** A 4000-fold decrease in the diffusion constant results in strong trapping at time t = 30 min with the gradient remaining pronounced even at time t=72 h.

We then varied one parameter at a time to determine how they individually affect antibody penetration after 30 min (Fig. 2b-e). Increasing Ab k_on_ by 100-fold from the baseline or decreasing Ab k_off_ by 100-fold abolished internal penetration (Fig.2b-c). This supports that proper Ab penetration is blocked when epitope binding affinity is high, matching our observations for the higher affinity anti-H3K9me2 Ab AM39239. Similarly, Ab penetrance into the nuclear interior could also be inhibited by increasing the Ag concentration (C_b_) by 50-fold or decreasing the diffusion coefficient (D_A_) by 50-fold (Fig. 2d-e).

We also investigated if failed penetration could arise when smaller changes are made to several variables simultaneously. Internal staining was prevented by simultaneously reducing Ab dissociation (k_off_) 10-fold while increasing Ab binding (k_on_) 10-fold and Ag concentration (C_b_) two-fold (Fig.2f). A similar result was achieved by a 13,(3)-fold reduction in the diffusion coefficient (D_A_) in combination with a 3-fold increase and 3-fold decrease in k_on_ and k_off_, respectively (Fig. 2g). Thus, a peripheral staining artifact can arise by synergistically modulating multiple parameters that impact Ab penetration.

A final parameter under experimental control is Ab incubation time. Simulations with our reference parameters showed that Abs progressively penetrate the nucleus over 30 min to ultimately produce a uniform distribution of signal (Extended Data Fig. 4b-c). Consistently, further extending the simulated incubation time to 72 h also resulted in a uniform signal even under conditions that previously produced exclusively peripheral staining (Fig.2h). Artifactual peripheral staining is thus predicted to be mitigated, in some cases, by increasing Ab incubation time. However, we note that extreme values for any of the parameters – or their combinations – could prevent this Ab penetration even at 72 h (Fig. 2i).

Taken together, our simulations argue that Ab performance in a closed volume is strongly influenced by multiple parameters, including epitope abundance, k_on_ and k_off_ that define Ab affinity, and diffusion rate. Combinations of low affinity, low epitope concentration and high diffusion consistently improve nuclear interior staining. In contrast, combinations of high affinity, high epitope concentration, and a low diffusion constant result in peripherally restricted staining of simulated nuclei. We termed this phenomenon Ab-trapping and next sought to experimentally test these computational predictions.

### High epitope abundance contributes to Ab-trapping

We began by testing whether high epitope concentration blocks Ab penetration. We did so by decreasing the abundance of H3K9me2 in three ways prior to detection with AM39239. First, we applied immunofluorescence to 3KO-MEFs which lack three of the five H3K9 methyltransferases - Suv39H1, Suv39H2, and SetDB1 - resulting in a 30% reduction of H3K9me2 levels^24^. As predicted, in the 3KO cells, the peripheral staining effect with AM39239 was substantially but not completely abrogated, and staining became more uniformly distributed throughout the nucleoplasm (Fig. 3a). Second, we applied a gentle trypsin digestion, which should remove histone tails like those marked with H3K9me2, in fixed HeLa cells. This limited digestion similarly promoted a more uniform labeling of chromatin in the nuclear interior (Fig. 3b). Finally, we performed immunofluorescence staining in tissue sections. Here, prolonged sample crosslinking and antigen retrieval by heating (see Methods) are expected to greatly reduce epitope abundance relative to standard immunofluorescence used for cultured cells. Matching this, AM39239 produced peripheral staining following standard immunofluorescence after short fixation and in the absence of antigen retrieval in cultured mouse embryonic limb fibroblasts and isolated retinal rod photoreceptors (Fig. 1c,d). However, this effect was entirely eliminated in cryosections of embryonic limb (Fig. 3c) and adult retina (Fig. 3d and Extended Data Fig. 5). Together, this supports that high epitope abundance is a major contributor to Ab-trapping for at least the anti-H3K9me2 Ab AM39239.

**Figure 3.**
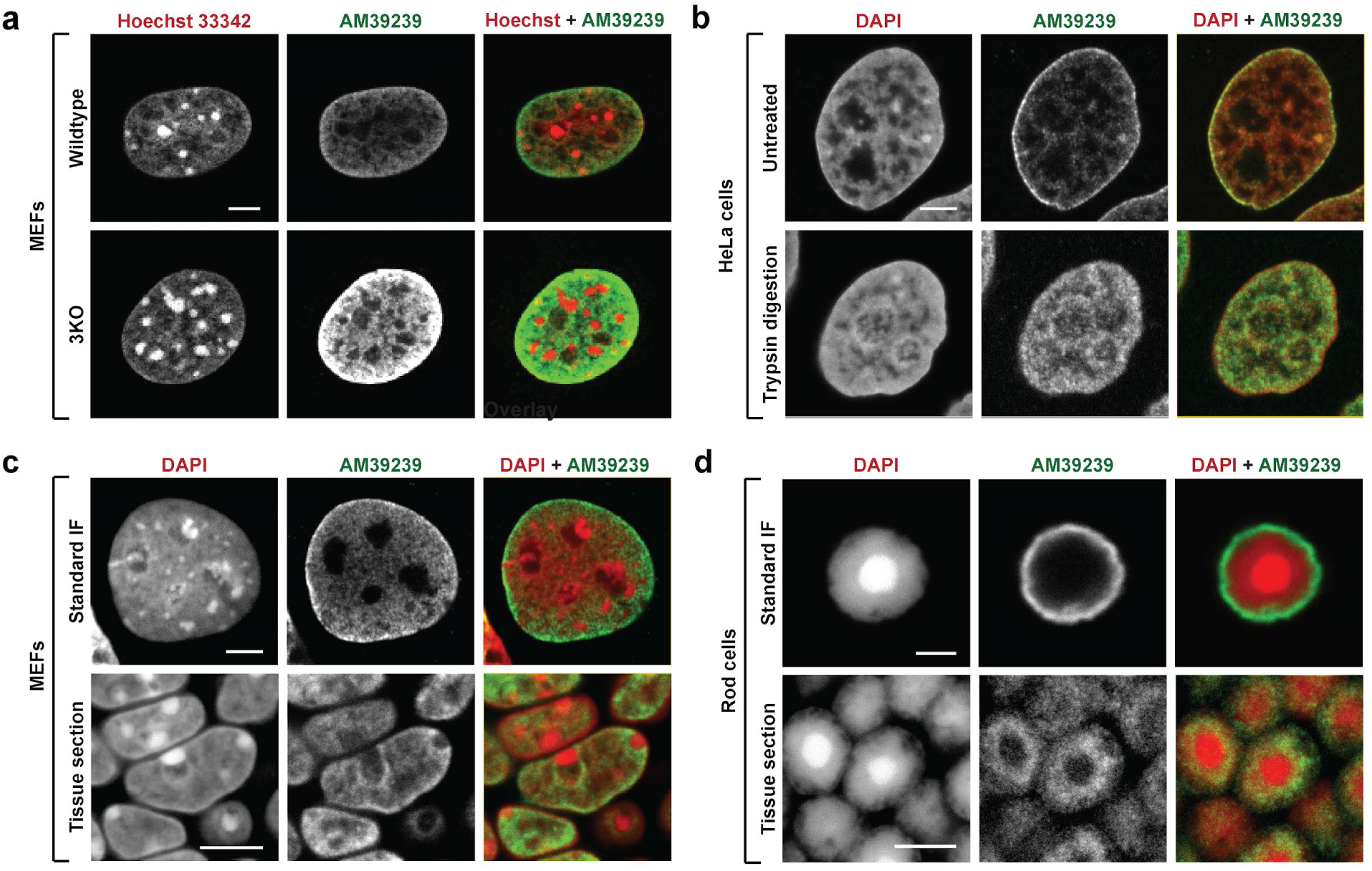
High epitope abundance contributes to Ab-trapping. **a**, Immunofluorescence staining with AM39239 in WT and 3KO MEFs. Ab signal in the nuclear interior is substantially increased in 3KO cells, although the gradient from the periphery towards the center is still present. **b**, Immunofluorescence staining with AM39239 in HeLa nuclei without (top) and with (bottom) nonspecific reduction of proteins by Trypsin digestion. Trypsin digestion notably enhances internal staining. **c,d**, Immunofluorescence staining with AM39239 in single cells (top layers) and in tissue cryosections (bottom layers). In cultured MEFs (c, top) and isolated rod photoreceptors (d, top), the nuclear staining is peripheral. In sections of embryonic limb (c, bottom), in addition to the peripheral rim, the Ab stains the nuclear interior of MEFs including the nucleolar periphery. In sections of retina (d, bottom), the Ab stains the internal heterochromatin layer. All images are single optical sections. Scale bars: 5 μm.

### High Ab affinity contributes to Ab-trapping

We further tested the prediction that Ab-trapping is linked to AM39239’s high affinity for H3K9me2, which is on average at least 6-fold greater than that of CMA317 (Extended Data Fig. 3). To do so, we examined the penetration dynamics of these two Abs by following progressive Ab staining in fixed cells over time using time-lapse microscopy. The Abs were preincubated with secondary fluorescently labeled Abs and mixed with Hoechst 33342, before incubating with fixed permeabilized cells. Strikingly, peripheral nuclear signal was initially observed for both Abs as well as even Hoechst staining. Full nuclear staining with CMA317 was then achieved after several hours, whereas staining with AM39239 was not completed even after 12 h (Extended Data Fig. 6 and Supplementary Video 1). Since both Abs target the same antigen, this confirms that the higher affinity of AM39239 promotes Ab-trapping.

### Increasing incubation time partially mitigates Ab-trapping

Our computational modeling and time-lapse experiments suggested that prolonged incubation with the primary Ab may improve Ab penetration (Fig. 2h-i and Extended Data Fig. 6). To test this, we used immunofluorescence to compare the performance of the three anti-H3K9me2 Abs after standard (1 h) and extended (48 h) incubation periods. For each Ab, both the intensity of staining and penetration into the nucleus improved with prolonged incubation (Fig. 4a). The enhancement in staining was particularly evident for AM39239. Specifically, although the chromatin in the interior was still weakly stained, in addition to the peripheral rim, the Ab now labeled the periphery of the nucleoli and nuclear channels formed by the invaginated nuclear envelope. This supports that Ab-trapping arises from the impaired diffusion of high-affinity Abs, a limitation that can be mitigated by extending staining times.

**Figure 4.**
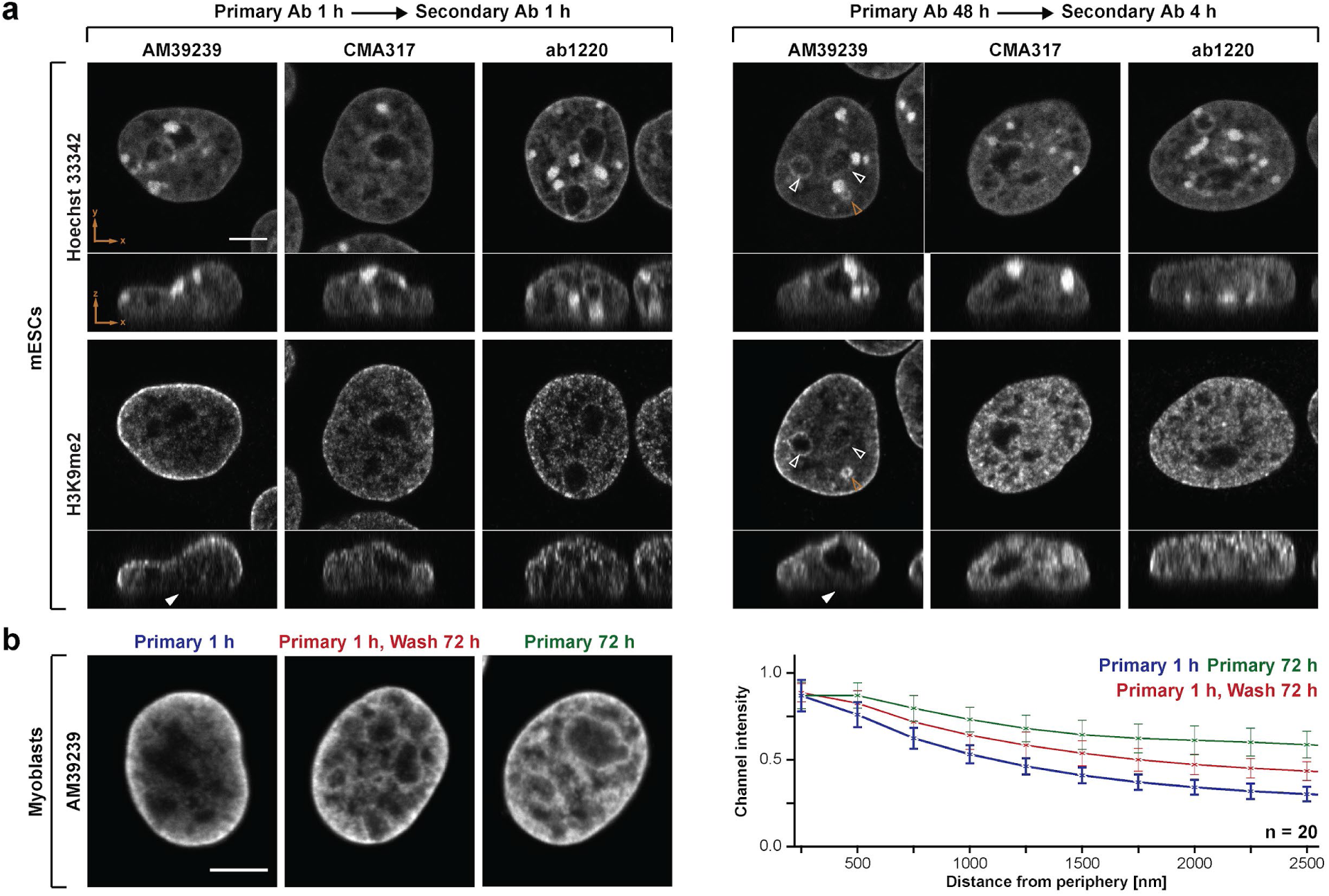
Increasing incubation time with primary Ab mitigates the Ab-trapping effect. **a**, Immunofluorescence staining with three anti-H3K9me2 Abs with standard (left) and extended (right) incubation times. AM39239 stains the nuclear periphery closest to the coverslip/slide only in the extended incubation (solid white arrowheads in XZ sections). Note the AM39239 staining at the nucleolar periphery (empty white arrowheads) and nuclear channels formed by the invaginated nuclear envelope (empty gold arrowheads) in extended incubations. **b**, Immunofluorescence staining with AM39239 (left) and quantification of radial signal distributions (right) in three conditions: (1) incubation with primary Ab for 1 h followed by 30 min wash, (2) incubation with primary Ab for 72 h followed by 30 min wash, and (3) incubation with primary Ab for 1 h followed by 72 h wash. The incubation with secondary Ab in all protocols was 1 h. All images are single optical sections. Scale bars: 5 µm.

We further reasoned that, if a high-affinity Ab has slow effective diffusion, then extending the washing time prior to applying the secondary Ab should also help counteract Ab-trapping. Indeed, increasing the washing time to 72 hours partially resolved Ab-trapping for AM39239, producing comparable internal nuclear staining as when the Ab is incubated on cells for an extended 72 h incubation (Fig. 4b). Thus, prolonged incubation is evidently sufficient for the IgG molecules to dissociate from the peripheral chromatin and diffuse into the nuclear interior. This also indicates that the quantity of the AM39239 Ab molecules that are initially trapped at the nuclear periphery is adequate to stain the entire nucleus without adding any additional new Ab molecules.

Other classical modifications of standard immunofluorescence, such as varying Ab concentration or adjusting blocking time, did not resolve artifactual AM39239 staining (Extended Data Fig. 7). Thus, increasing Ab incubation time emerges as the most reliable and easily implementable adjustment within existing staining protocols to reduce Ab-trapping.

### Ab-trapping impacts genomic assays

We now searched for other experimental contexts where Ab-trapping could produce artifactual results. We focused on the Ab-based genomics technology CUT&Tag^9^ where Ab-trapping could also create excessively peripheral signal for anti-H3K9me2 Abs. Specifically, we examined inverted rod cells and detected peripherally positioned chromatin found in lamina-associated domains (LADs) with anti-lamin B1 (Abcam, ab16048), euchromatin with anti-H3K9ac (Active Motif, AM39685), and H3K9me2 with the three anti-H3K9me2 Abs.

As expected, rods displayed a characteristically inverted genome organization with LADs coinciding with H3K9ac-marked euchromatin (Fig. 5a,b). As such, the euchromatic LADs of rods anti-correlate with the heterochromatic LADs of conventionally organised ESCs. However, as with our previous immunofluorescence experiments, specific anti-H3K9me2 Abs produce distributions of signal consistent with Ab-trapping in rod cells. In particular, ab1220 signal correctly coincides with heterochromatic non-LADs in the nuclear interior. By contrast, both AM39239 and CMA317 signals are incorrectly present at euchromatic LADs at the periphery. These results were mirrored by imaging after staining non-fixed rods in conditions and buffers from the CUT&Tag protocol (Fig. 5c) and quantifying radial signal distribution (Fig. 5d).

**Figure 5.**
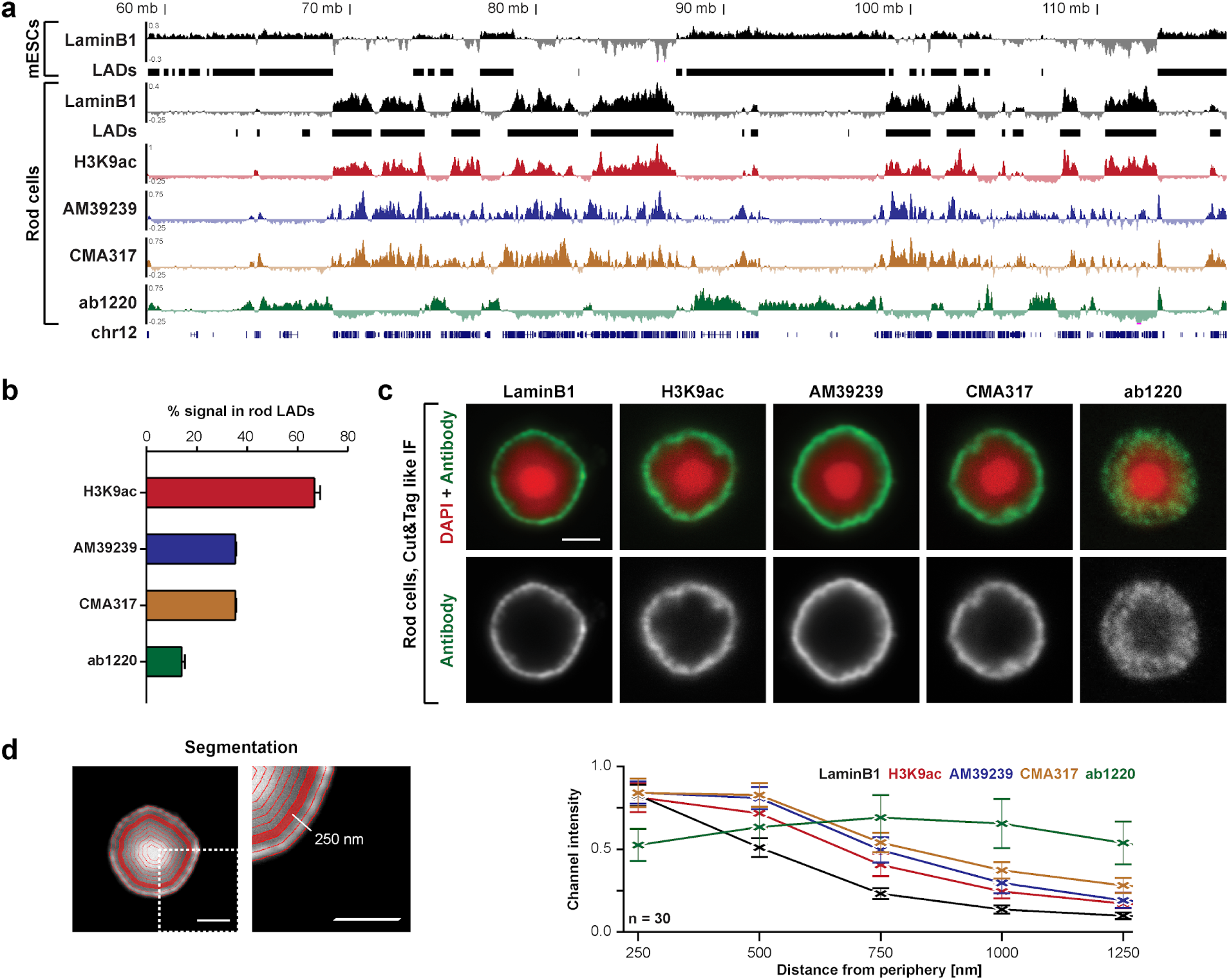
Anti-H3K9me2 Abs display peripheral Ab-trapping in CUT&Tag. **a**, CUT&Tag tracks in G4 mESCs and rod photoreceptors for indicated Abs with Ref-seq genes shown below. LADs called from LMNB1 CUT&Tag are shown below. In all cases, CUT&Tag tracks show log_2_ signal after subtraction of IgG control. **b**, Corresponding fraction of Ab CUT&Tag signal that overlaps with rod cell LADs. **c**, Matching immunofluorescence staining performed under conditions analogous to CUT&Tag in isolated rod cells. **d**, Schematics of nuclear segmentation (left) and quantification of radial signal distributions (right) of the indicated Abs. All images are single optical sections. Scale bar: 2 µm.

For AM39239, this Ab-trapping in rods matches our previous microcopy stainings (Fig. 1e) and published CUT&RUN data^21^. However, we note that CMA317 staining differed depending on the experimental methodology. Though showing peripheral Ab-trapping under CUT&Tag conditions (Fig. 5c), CMA317 correctly labeled chromatin throughout the nucleus in our previous standard immunofluorescence (Fig. 1e). Together, this indicates that Ab-trapping can impact both microscopy and genomics assays, but in a variable manner influenced by exact experimental conditions.

### Ab-trapping affects multiple Abs, epitopes and structures

Our modeling suggests that Ab-trapping can in principle affect any Ab or cellular structure with epitopes that are confined to a limited volume. We thus examined several alternative scenarios where artifactual peripheral staining could emerge in a manner dependent on Ab affinity and/or epitope abundance.

We initially applied standard immunofluorescence to a HeLa cell line that heterogeneously expresses H2A-GFP at variable levels across different cells. Strikingly, the anti-GFP Ab A-11122 artifactually stained the nuclear periphery in cells with high H2A-GFP expression. By contrast, lower expressing cells with a reduced GFP epitope abundance were unaffected (Fig. 6a). In parallel, cells were also co-stained with an anti-GFP nanobody that binds GFP with a single monomeric variable domain and is thus ten-fold smaller than rabbit IgGs^30^. Accordingly, the nanobody stained GFP throughout the entire nucleoplasm independently of GFP expression level, presumably due to their lower k_on_ and/or higher diffusion coefficient (D_A_) (Extended Data Fig. 8 a,b). Based on our modeling, this was an expected result. Simulations using the published parameters for both antibody and nanobody revealed that the anti-GFP rabbit Ab produces strong peripheral staining at high GFP expression (Extended Data Fig. 8c,d). However, the simulated nanobody yields almost homogeneous nuclear staining with a weak gradient. Thus, even GFP Abs are subject to Ab-trapping when under the synergistic effects of high epitope abundance, high Ab affinity and slow diffusion.

**Figure 6.**
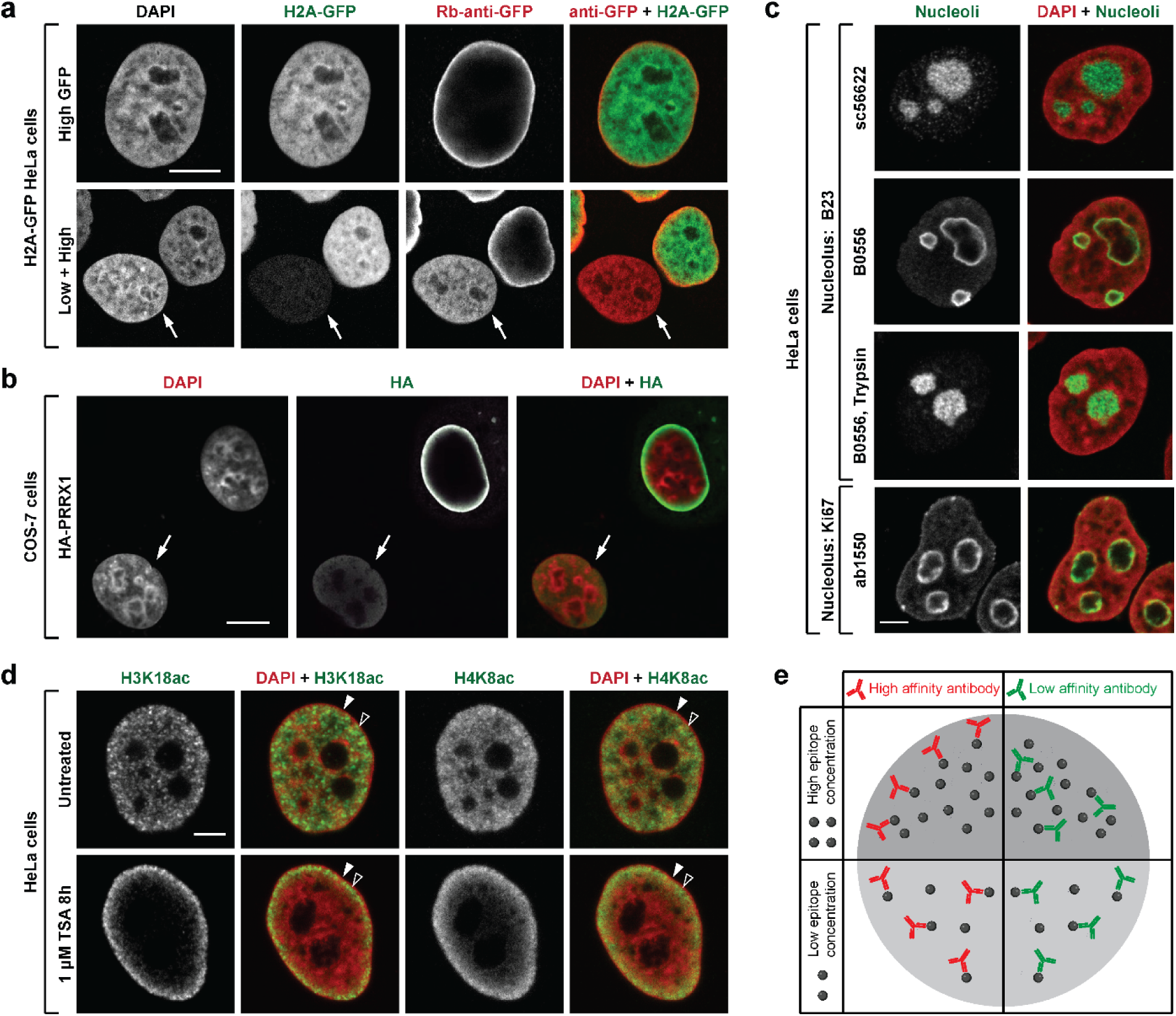
Multiple Abs and target epitopes exhibit the trapping effect. **a**, Immunofluorescence staining with anti-GFP Ab in nuclei of HeLa cells with variable expression of H2A-GFP. Peripheral Ab-trapping is observed in highly expressing H2A-GFP cells but not in cells with lower expression (arrow). **b**, Immunofluorescence staining with anti-HA Ab in nuclei of COS-7 cells expressing HA-tagged transcription factor PRRX1. Note the false nuclear peripheral staining in highly expressing cells and nucleoplasm staining in cells with lower expression (arrow). **c**, Immunofluorescence staining with anti-B23 (top) and anti-K67 (bottom) Abs in HeLa cells without and with nonspecific reduction of proteins by Trypsin digestion (third row). Note that the false B23 immunostaining can be avoided by using a different Ab (sc-56622) or mild protein digestion. **d**, Immunofluorescence staining with anti-H3K18ac or anti-H4K8ac Abs in HeLa cells without (top) and with increased histone acetylation via Trichostatin A (TSA) treatment (bottom). Homogeneous staining with H3K18ac and H4K8ac Abs throughout the internal euchromatin becomes peripheral after the increase in epitope abundance. Note that in contrast to AM39239 Ab-trapping at the extreme nuclear periphery (Fig.1), euchromatin Ab-trapping (empty arrowheads) occurs more internally than the peripheral rim of heterochromatin (solid arrowheads). All images are single optical sections. Scale bars: a-b 10 µm, c-d 5 µm. **e**, Schematic illustrating the synergistic effect of high Ab affinity and high epitope concentration. Schematic created with BioRender.com.

We further examined other tagged proteins for Ab-trapping. Staining the HA-tagged transcription factor PRRX1 results in a nucleoplasm-wide signal when its expression is low but only false peripheral staining when its expression is high (Fig. 6b). Likewise, a previous study demonstrated that anti-GFP or anti-FLAG Abs display Ab-trapping at the nucleolar surface when fibrillarin-GFP and EGFP-FLAG-B23 are overexpressed in HeLa cells^31,32^. Matching this, we found that the abundant endogenous B23 protein also displays Ab-trapping depending on the Ab used. Specifically, the sc-56622 Ab stains nucleoli uniformly, whereas the alternative Sigma B0556 Ab stains only the nucleolar periphery causing a typical Ab-trapping effect that is abrogated after mild digestion (Fig. 6c). Similarly, staining of another abundant intra-nucleolar protein Ki67 with Abcam ab15580 was also restricted to the nucleolar periphery (Fig. 6c). Thus Ab-trapping can affect multiple tagged and endogenous proteins in distinct cellular structures.

We finally tested for potential Ab-trapping in other abundant histone modifications that define euchromatin instead. Under normal conditions Abs against H4K8ac or H3K18ac displayed enriched signals in the nuclear interior in HeLa cells (Fig. 6d). However, elevating the concentration of acetylated histones in HeLa cells following treatment with the HDAC inhibitor Trichostatin A, increased labeling to an Ab-trapping-like sub-peripheral rim.

In summary, Ab-trapping can affect Abs targeting a wide range of epitopes, including both endogenous and ectopically expressed proteins within the nucleus and nucleolus.

## DISCUSSION

Our study presents a detailed characterization of a common yet previously undescribed immunolabeling artifact - Ab-trapping. Here, by using modeling and experimental validation, we identified a combination of parameters that drive this artifact. These include (i) a high epitope concentration confined to a limited volume and (ii) high Ab affinity that together restrict Ab penetration (Fig. 6e). We further demonstrated the trapping phenomenon for Abs targeting a variety of endogenous and ectopically expressed proteins, emphasizing its widespread prevalence.

The resulting restriction to Ab binding poses a significant challenge in any assay relying on Ab penetration into densely packed nuclear volumes, including immunofluorescence labeling and CUT&Tag sequencing. While we describe Ab-trapping for nuclear and nucleolar epitopes, other abundant cellular features likely present similar issues. Critically, many emerging technologies require the *in situ* or *in vivo* application of Abs. This includes recent single-cell epigenomics methods to map chromatin features in individual nuclei and technologies that spatially resolve the distribution of proteins in tissues^9,11–14^. Ab-trapping can thus have important consequences, which the scientific community must be aware of going forward.

The phenomenon of Ab-trapping is known to many microscopists. As such, peripheral staining patterns of the nucleus have been successfully recognized as artifactual in some cases (e.g., personal communications of Evgenya Popova, Sergei Grigoryev, Robert Schneider and others), including in the nucleolus as described above^31,32^. However, there are also other examples where artifactual peripheral staining has been erroneously interpreted as a true result. This includes the artifactual near-exclusive peripheral staining of H3K9me2 and/or H3S10ph in interphase nuclei and condensed mitotic chromosomes^33,34^, as well as the artifactual H3K9me2 peripheral euchromatic signal in mouse rod photoreceptor cells^21^. Here, we demonstrate these observations result from Ab-trapping.

Our experiments showed that one of the three anti-H3K9me2 Abs, ab1220, functions in both CUT&Tag and immunostaining. By contrast, the CMA317 Ab does not cause Ab-trapping in standard immunostaining but does under CUT&Tag conditions, which is likely related to applying Ab to unfixed cells and/or within a specific buffer. Conversely, the Active Motif AM39239, produces artifactual results in CUT&Tag and standard immunostaining of single cells but not in tissue sections with reduced epitopes. This variability highlights the care that must be taken when interpreting results from even the same Ab across methodologies.

While Ab-trapping is anecdotally known in the microscopy community, a model of artifactual peripheral staining at the cellular scale has been lacking until now. By making use of available biophysical measurements, our model captures key aspects of antibody staining kinetics, and future work can expand on the simplifying assumptions made here. First, cell nuclei are not perfectly spherical, so artifacts arising from the staining process could be influenced by target geometry. Second, we do not model fluid advection or non-specific binding events, both of which could affect staining dynamics. Third, considerable uncertainty remains regarding reference parameters, including Ab affinity and diffusion coefficients. Finally, we did not model bivalent or cooperative binding. Incorporating such details may enable computational screening to avoid Ab-trapping and other artifacts across a broad range of staining scenarios.

We would like to alert the scientific community that there are many other commercially available Abs that target nuclear epitopes and display apparent Ab-trapping on the manufacturer’s websites. These are exemplified by Abs from ThermoFisher Scientific for H1 and H4 acetylation, Active Motif for histone H3, and Abcam for H3K27ac (Supplementary Table 1). Importantly, this does not imply that these Abs lack specificity. Rather, their high affinity and/or the abundance of their target epitopes likely predisposes them to Ab-trapping. Consequently, such Abs can still produce reliable results if appropriate measures to mitigate Ab-trapping are employed. To assist researchers in recognizing and mitigating Ab-trapping, we recommend following straightforward best practices:

**(1) Peripheral Staining Caution.** Immunostaining that is confined to the periphery of the nucleus, nucleolus or similar confined structures should be interpreted with caution. If possible, use a different Ab (or tagged protein expression) to confirm the peripheral localization.
**(2) Expression level variability.** When immunodetecting ectopically expressed proteins via protein tags, ensure that a sufficient number of cells are analyzed covering a wide range of expression levels. If a peripheral signal is observed in only a fraction of highly expressing cells, it is likely the result of Ab-trapping caused by high epitope abundance.
**(3) Optimization of staining conditions.** When using an untested primary Ab, trial different staining and washing times (from 1 to 72 h) to optimize conditions. Likewise, various incubation conditions might influence Ab-trapping, e.g., differential performance of CMA317 in CUT&Tag and PBS buffers.
**(4) Confirmation by using cryosections.** Whenever possible, validate staining patterns observed in isolated or cultured cells by using tissue cryosections where longer crosslinking and antigen retrieval reduces epitope abundance.

## EXTENDED DATA FIGURES AND LEGENDS

**Extended Data Figure 1.**
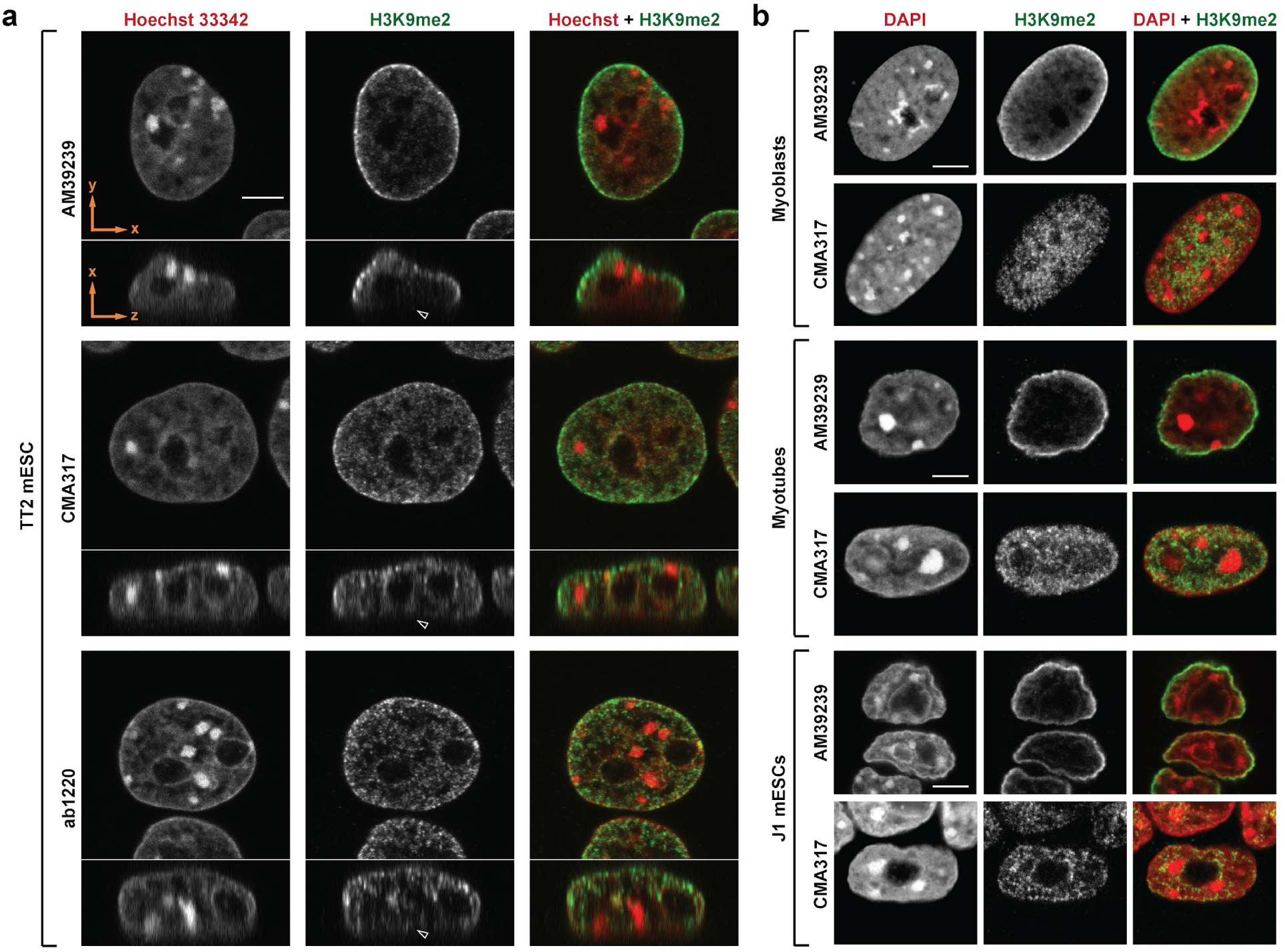
Immunofluorescence staining using three different anti-H3K9me2 Abs in multiple cell-types. **a**, Immunofluorescence staining with single anti-H3K9me2 Abs in TT2 mESCs nuclei. XY (top) and XZ (bottom) views are shown. Arrowheads in XZ sections point at the nuclear periphery adjacent to the substrate, which lacks staining with AM39239. **b**, Immunofluorescence staining with pairs of distinct anti-H3K9me2 Abs in nuclei of cultured mouse myoblasts, myotubes (differentiated from myoblasts) and J1 mESCs. All images are single optical sections. Scale bars: 5 μm.

**Extended Data Figure 2.**
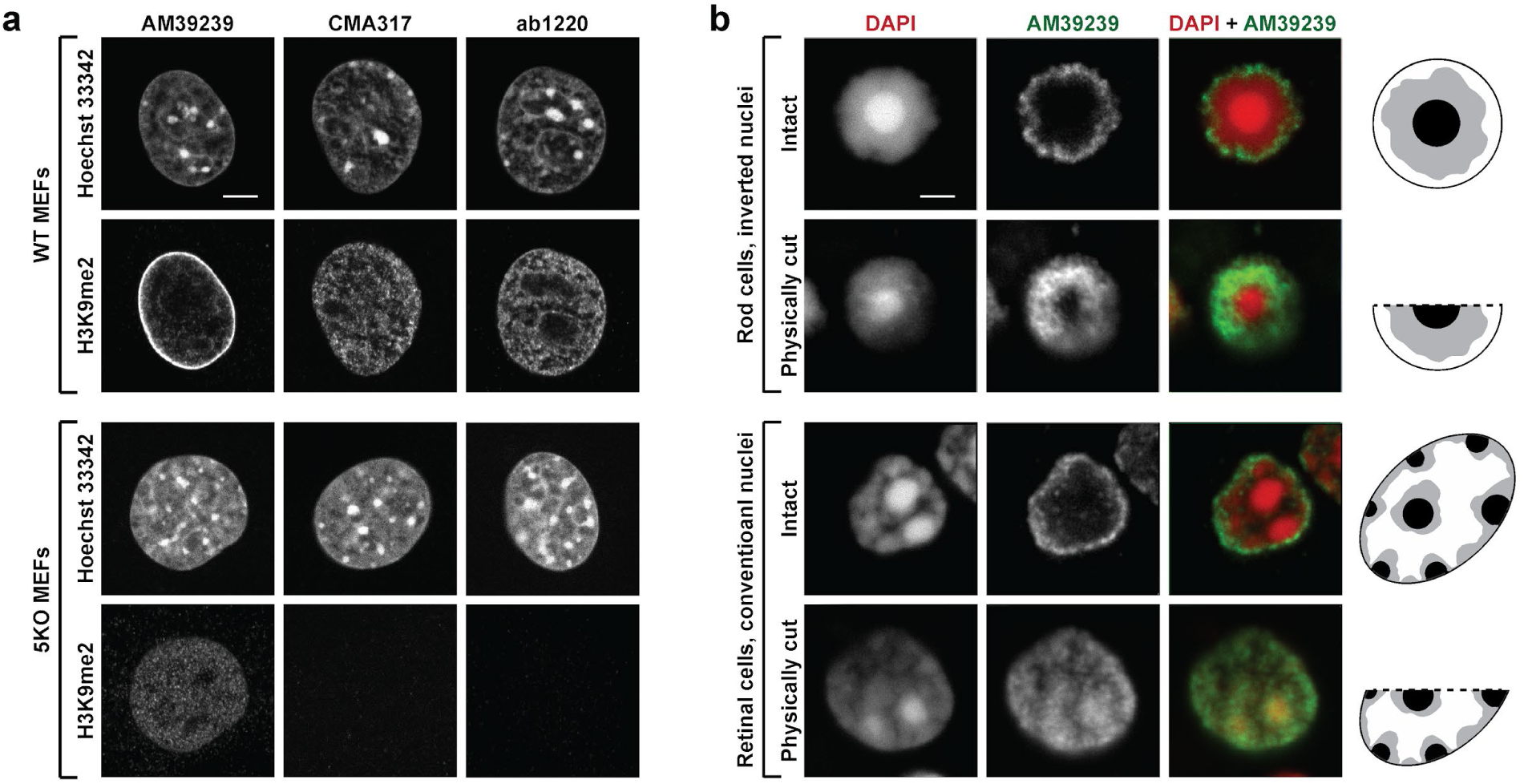
Tests of Ab specificity and penetration capacity. **a**, Immunofluorescence staining with single anti-H3K9me2 Abs in WT (top) and 5KO MEFs nuclei lacking H3K9me2 (bottom). Note the absence of CMA317 and ab1220 signals in 5KO-MEF nuclei, while AM39239 produces homogeneous residual staining throughout the whole nucleus. **b**, Immunofluorescence staining with AM39239 in nuclei of retinal cells fixed in suspension and physically cut within cryosections. Intact rod and non-rod nuclei within the 14 µm thick sections display staining of the peripheral rim typical for the Ab (top layers). Cells at the cryosection surface with physically cut nuclei display chromatin staining throughout the cut surface (bottom layers). All images are single optical sections. Scale bars: 5 μm.

**Extended Data Figure 3.**
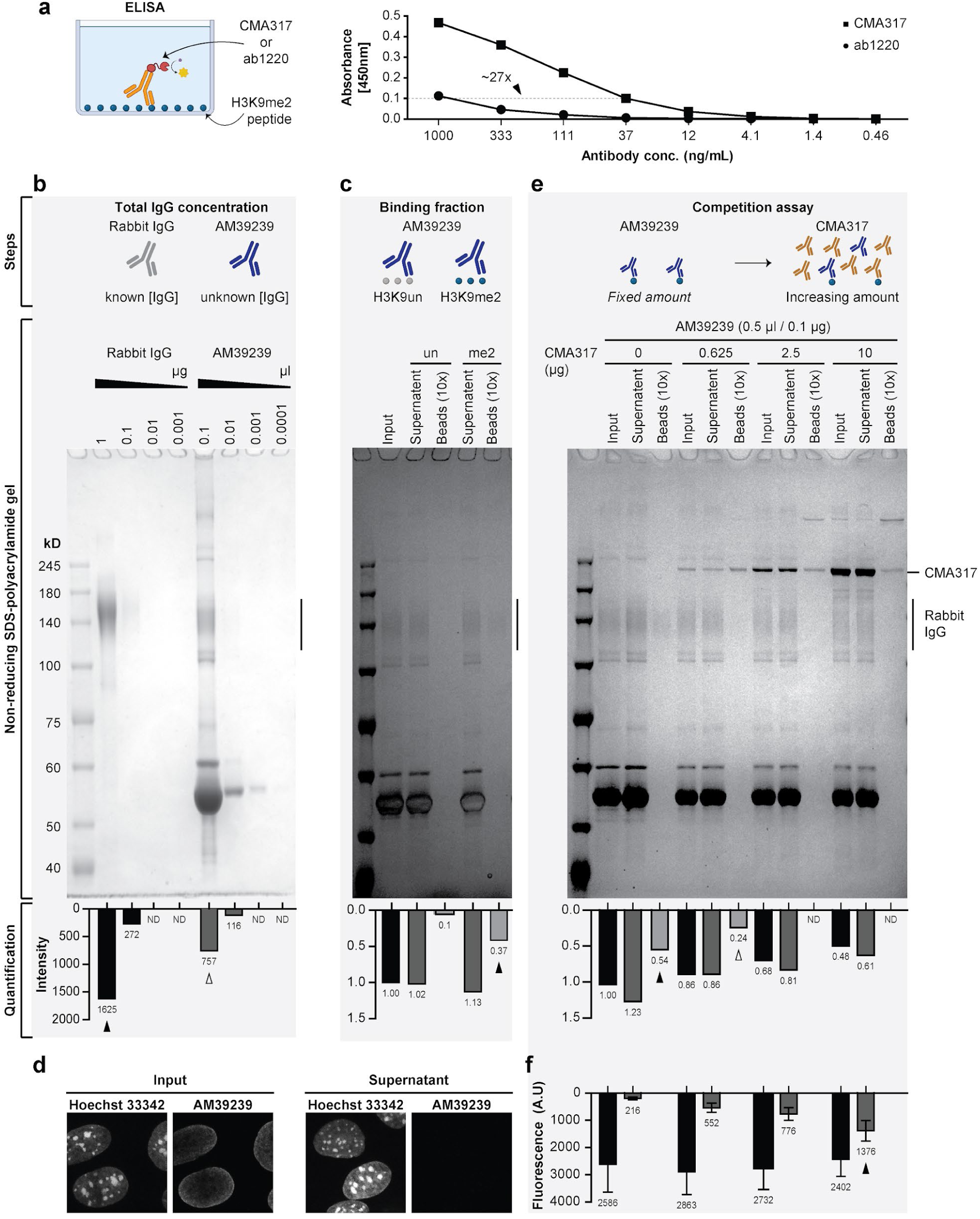
Estimating the relative binding affinities of anti-H3K9me2 Abs. **a**, ELISA profiling of the monoclonal CMA317 and ab1220 Abs comparing the binding affinities to the H3K9me2 epitope. H3K9me2 peptide-coated wells were incubated with a three-fold dilution series of Abs. Similar signals were observed at 27-fold different dilutions between the two Abs (e.g., 1,000 ng/ml CMA317 and 37 ng/ml ab1220) (arrowhead). **b**, Measuring the total concentration of IgG molecules in AM39239 anti-sera. The strategy is schematically illustrated (top). Known amounts of purified rabbit IgG and a serial dilution series of AM39239 were loaded onto a non-reducing SDS-polyacrylamide gel (middle) and the intensities were measured (bottom). 0.1 μl of AM39239 (empty arrowhead) contains ∼0.5 μg IgG (arrowhead), indicating an IgG concentration of ∼5 mg/ml in AM39239. **c**, Measuring the total H3K9me2 binding fraction of AM39239 anti-sera. The strategy is schematically illustrated (top). AM39239 IgG captured using H3K9 or H3K9me2 beads were loaded onto an SDS-polyacrylamide gel (middle) and the intensities were measured (bottom). The H3K9me2-bound fraction represents ∼4% (arrowhead) of total IgG (∼0.2 mg/ml) in AM39239. **d**, AM39239 immunofluorescence staining of input (left) and supernatant post-binding to the H3K9me2 beads (right) as represented in c. Lack of signal after binding indicates that most of the specific anti-H3K9me2 IgG molecules were captured by the beads. **e**, Measuring relative AM39239 binding affinity to CMA317 via competition assay. The strategy is schematically illustrated (top). AM39239 (0.1 μg of the H3K9me2-bound fraction) and CMA317 (0-10 μg) were mixed, captured using H3K9me2 beads and loaded onto an SDS-polyacrylamide gel (middle) and the intensities were measured (bottom). A 6-fold excess of CMA317 (empty arrowhead) reduced the capture of AM39239 (arrowhead) by roughly 50%. **f**, Quantification from AM39239 immunofluorescence staining using the input and bead supernatant fractions as represented in e. Signal was significantly reduced in the supernatant compared to input samples, even at 100-fold excess of CMA317 (arrowhead). AM39239 thus contains a small fraction of very high-affinity IgG molecules that contribute to the immunofluorescence signal. Schematics created with BioRender.com.

**Extended Data Figure 4.**
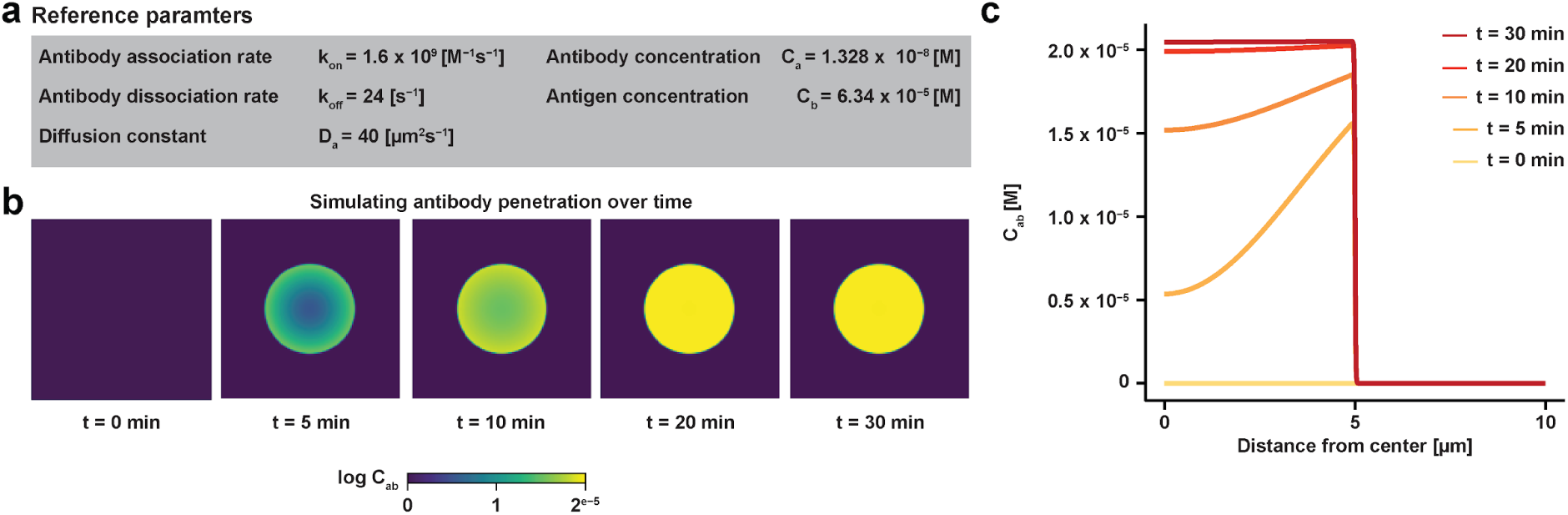
Reference parameters and additional perturbations of Ab staining simulations. **a**, Reference parameters for simulations. **b**, Time evolution of the concentration field C_AB_ with the reference parameters listed in (a). Over time, a concentration gradient develops and dissipates within approximately 20 min. Note the different color scale from the main text. **c**, Concentration field C_AB_ as a function of the distance from the center of the nucleus over time. Initially, a concentration gradient forms as the antibody diffuses inward and binds to the antigen. Over time, this gradient progressively diminishes as the system reaches equilibrium.

**Extended Data Figure 5.**
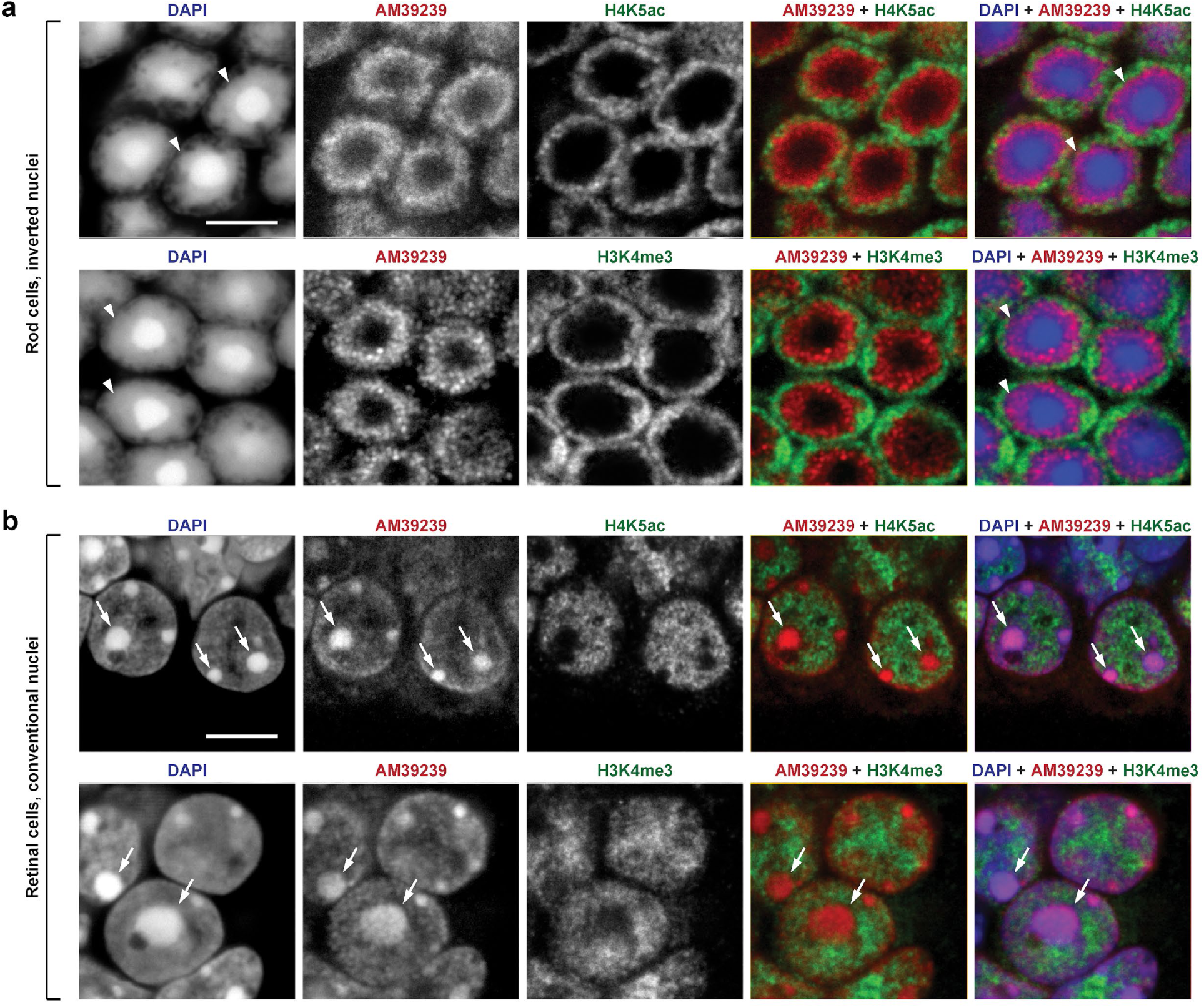
Immunofluorescence staining of rods and non-rod cells in retinal sections. **a-b**, Immunofluorescence staining with pairs of AM39239 and euchromatic (H4K5ac or H3K4me3) Abs in retinal sections. Nuclei of rods (a) or non-rod neurons (b) are shown. Note that in rods (a) AM39239 stains the internal layer of heterochromatin surrounding the central chromocenter (arrowheads), while in non-rods (b) heterochromatin stained with AM39239 (red, arrows) is mostly excluded from euchromatin (green). All images are single optical sections. Scale bars: 5 μm.

**Extended Data Figure 6.**
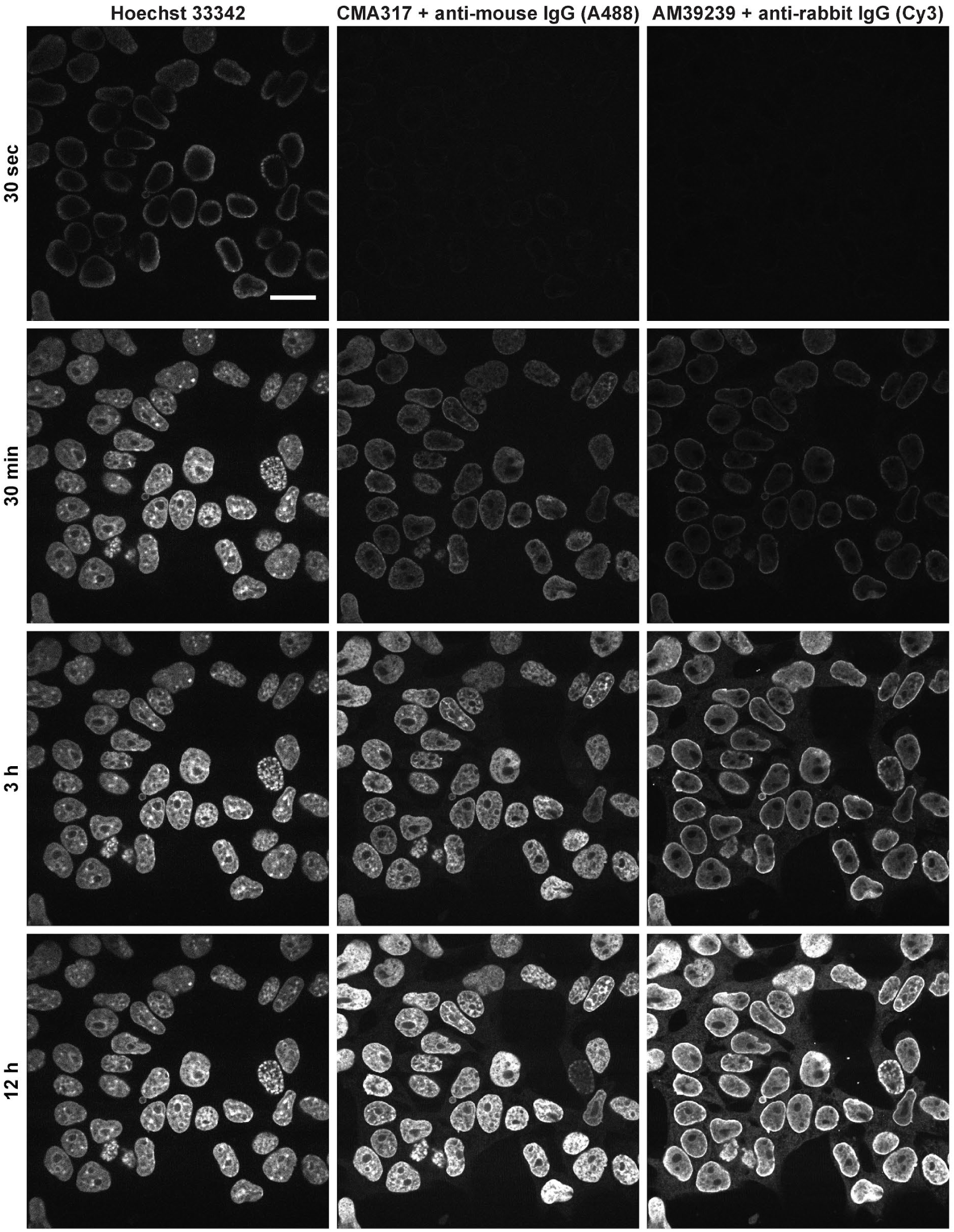
Time-lapse imaging of simultaneous immunofluorescence staining with CMA317 and AM39239 Abs. Immunofluorescence staining in nuclei of fixed and permeabilized HeLa cells incubated with CMA317 (preincubated with Alexa Fluor 488-conjugated anti-mouse IgG), AM39239 (preincubated with Cy3-conjugated anti-rabbit IgG), and Hoechst 33342. Staining dynamics were captured with time-lapse imaging for 16.5 hours with 5 min intervals. Note that nuclear rim staining is observed for all probes at early time points. Nuclear interior regions gradually become stained with Hoechst and CMA317, whereas the peripheral staining with AM39239 persists even after 12 h of incubation. See **Supplemental Video 1** for the full time-series. Scale bar: 20 µm.

**Extended Data Figure 7.**
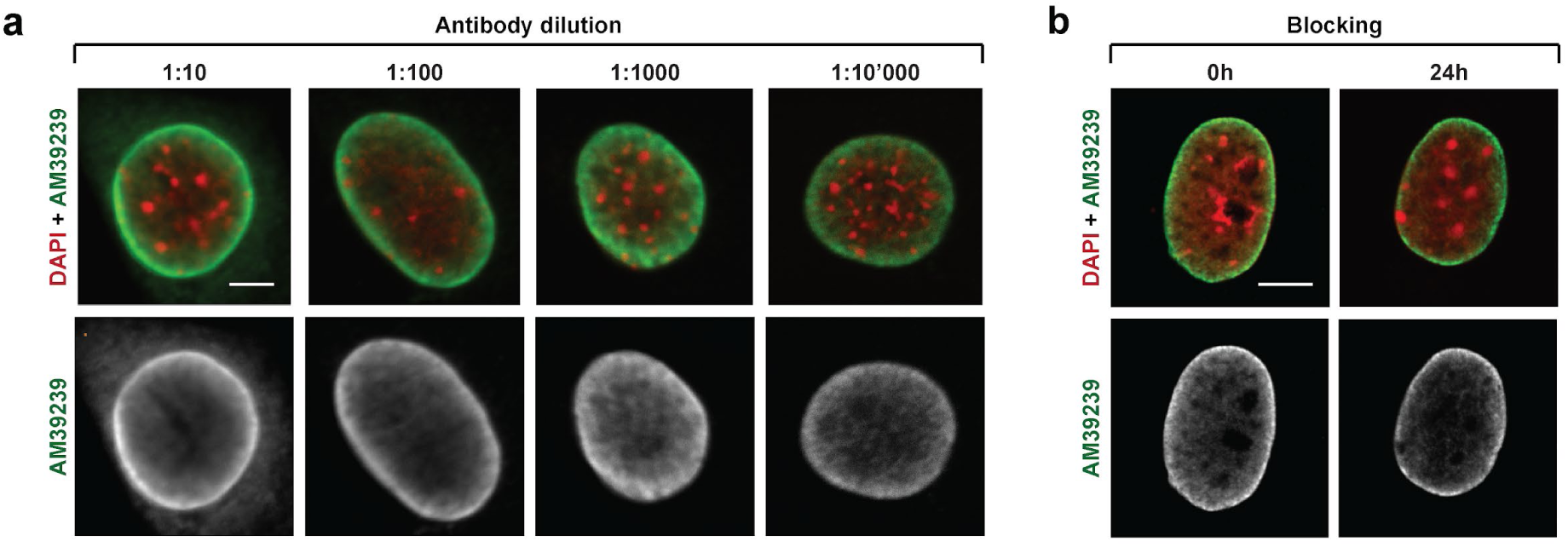
Conventional changes in immunofluorescence staining protocol do not mitigate peripheral staining with AM39239 Ab in myoblasts. **a-b**, Immunofluorescence staining with AM39239 in nuclei of myoblasts with varying concentration of Ab (a) or with and without blocking (b). Neither altered primary Ab concentration nor prolonged blocking changed the staining pattern. All images are single optical sections. Scale bars: 5 µm.

**Extended Data Figure 8.**
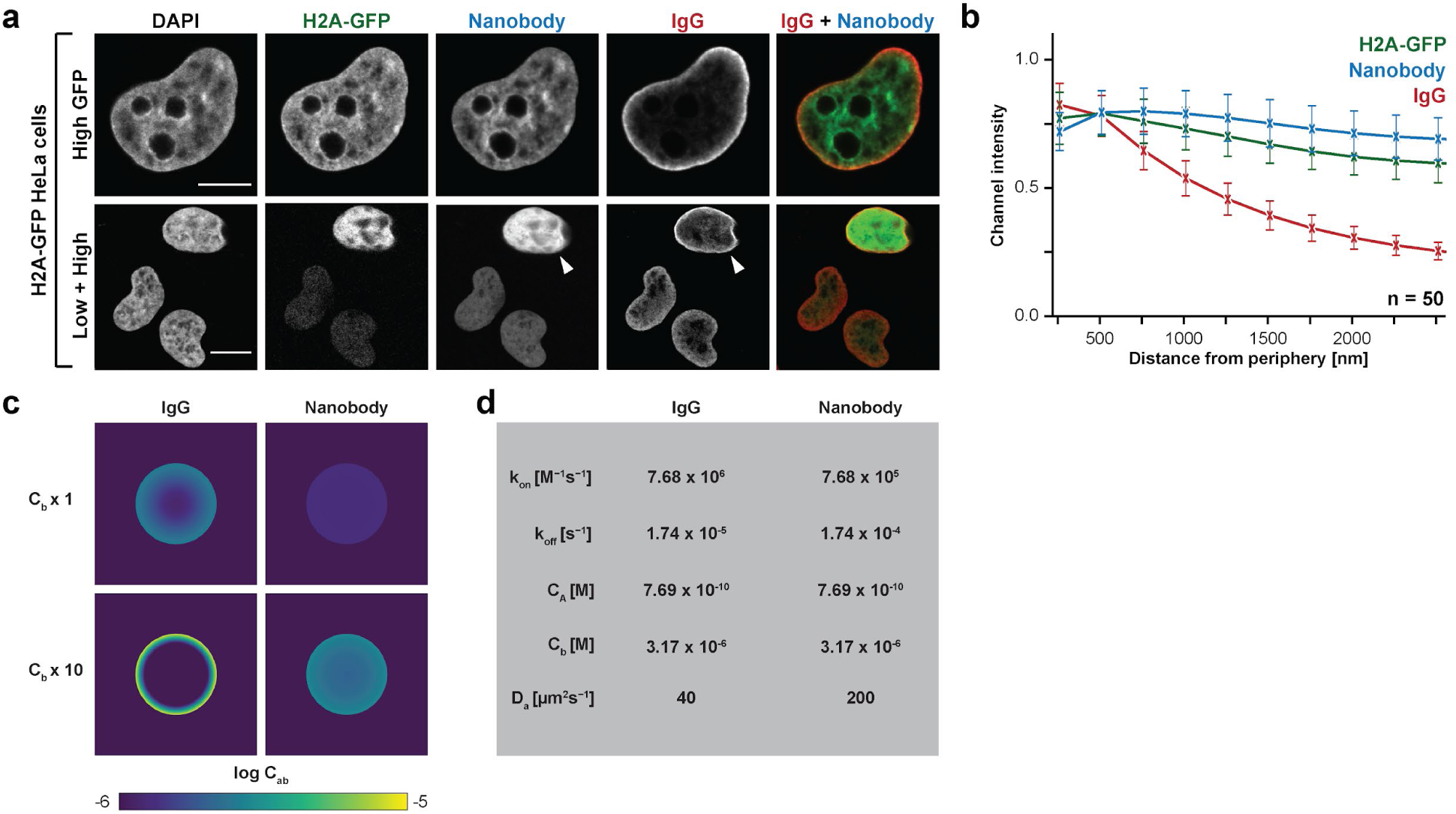
Anti-GFP nanobodies are not affected by Ab-trapping. **a**, Immunofluorescence staining with anti-GFP rabbit IgG and nanobody in nuclei of HeLa cells with variable expression of H2A-GFP. Note that the anti-GFP IgG is trapped at the nuclear periphery in cells highly expressing H2A-GFP (arrowhead), while the anti-GFP nanobody stains the whole nucleoplasm. All images are single optical sections. Scale bars: 10 µm. **b**, Quantification of radial signal distributions of indicated Abs confirms peripheral enrichment of IgG. **c-d**, The concentration field C_AB_ for a simulated anti-GFP nanobody or IgG at incubation time t=30 min (c) with corresponding simulation parameters (d). Simulation recapitulates the difference in staining pattern between IgG and nanobody in highly expressing cells. Note the different color scale from the main text.

## METHODS

### Cell culture

Mouse TT2 ESCs and immortalized embryonic fibroblasts (iMEFs) were obtained from Yoichi Shinkai^24^. TT2 ESCs were cultured in DMEM supplemented with 10% fetal calf serum (FCS), 1x nonessential amino acids, sodium pyruvate, 2–mercaptoethanol, 1x penicillin/streptomycin, and 1000 U/ml LIF (Nacalai Tesque). iMEFs, TKO (Setdb1, Suv39h1/2 triple knockout) and 5KO (Setdb1, Suv39h1/2, G9a, Glp quintuple knockout) iMEFs were maintained in the same medium as TT2 ESCs but without LIF.

The mouse myoblast cell line Pmi28 was grown in F-10 Nutrient Mixture (Ham) supplemented with 20% FCS and 1% penicillin/streptomycin. Naive J1 mESCs, MEFs and H2A-HeLa cells were obtained from the Leonhardt laboratory. Both MEFs and HeLa cells were cultured in DMEM supplemented with 10% FCS and 1% penicillin/streptomycin. J1 mESCs were cultured in serum-free media consisting of 50% N2B27, 50% DMEM/F12, 2i, 1000 U/ml recombinant LIF, 0.3% BSA, 2 mM L -glutamine and 100 U/ml penicillin. All cultures were maintained at 37°C and 5% CO₂ and regularly checked for mycoplasma contamination.

Mouse G4 embryonic stem cells (ESCs) (XY, 129S6/SvEvTac x C57BL/6Ncr F1 hybrid) were cultured as previously described^35,36^ on mitomycin–inactivated CD1 mouse embryonic fibroblast (MEF) feeders in gelatinized dishes at 37°C, 7.5% CO₂. ESCs were maintained in knockout DMEM (Gibco, 10829018) containing 4.5 mg/ml glucose and sodium pyruvate, supplemented with 15% FCS (PAN-Seratech, 2602P122011), 10 mM glutamine (Biochrom, K0302), 1x penicillin/streptomycin (Biochrom, A2213B), 1x nonessential amino acids (Gibco, 11140050), 1x nucleosides (Sigma, ES008D) , 0.1 mM β-mercaptoethanol (Sigma, 805740), and 1000 U/ml leukemia inhibitory factor (LIF) (Millipore, ESG1106). Medium was changed daily, and cells were passaged every 2–3 days or cryopreserved at 1 × 10⁶ cells per vial in ESC medium containing 20% FCS and 10% DMSO (Sigma, D2650). Both ESCs and feeder cells were routinely tested for Mycoplasma contamination (Lonza, MycoAlert detection kit).

The COS-7 cells were cultured and transiently transfected with a plasmid encoding HA-tagged PRRX1 (S104G variant) under the control of a CMV promoter as described before^37^.

### Animal husbandry

B6.Cg-Tg(Nrl-EGFP)1Asw/J (NRL-GFP) mice^38^ were derived by in vitro fertilization using cryopreserved sperm into the C57Bl.6/J strain. The line was maintained on a C57Bl.6/J background and used to isolate retinal rod cells. Other mouse samples were collected from the C57Bl.6/J strain sourced from Inotiv. All mice were housed in a centrally controlled environment with a 12 h light and 12 h dark cycle, temperature of 20-22.2°C, and humidity of 30–50%. All mice had access to food and water *ad libitum*. Bedding, food and water were routinely changed. All experiments involving animals were carried out following institutional guidelines as approved by LaGeSo Berlin and following the Directive 2010/63/EU of the European Parliament on the protection of animals used for scientific purposes.

### Tissue sampling

Fixation of tissues and preparation of cryosections was done as previously described^39^. Briefly, tissues were fixed with 4% formaldehyde in PBS for 20–24 h, washed with PBS, incubated in sucrose with increasing concentrations (10%, 20% and 30%) and transferred into embedding molds (Peel-A-Way Disposable Embedding Molds, Polysciences Inc., USA) filled with Jung freezing medium (Leica Microsystems). Tissue cryoblocks were frozen by immersing the molds into a −80°C ethanol bath and stored at −80°C. Cryosections of 16-20 μm thickness were cut using a Leica Cryostat (Leica Microsystems), collected on SuperFrost microscopic slides (SuperFrost Ultra Plus, Roth, Germany), immediately frozen, and stored at −80°C before use.

A retinal cell suspension for physical bisectioning was generated from four retinas of C57Bl.6/J adult mice. Retinas were dissociated using Papain Dissociation System (Worthington, #130-094-802) as described previously^40^. Briefly, four retinas from two mice were dissociated in papain solution for 60 min at 37°C, while shaking at 700 rpm. Afterwards, retinas were triturated with a 1 ml micropipette and transferred to a new tube containing a mix of EBSS, DNAse and albumin-ovomucoid inhibitor. Cell suspensions were triturated again with a glass Pasteur pipette up to ten times until tissue fragments were no longer visible. Finally, 250 µl of EBSS/albumin-ovomucoid inhibitor was added, and the single cell suspension was cleared of cell clumps by filtration (pluriStrainer Mini 70 μm). 1:1 ratio of 10% BSA/PBS was added to the final single cell suspension to form a cushion for centrifugation at 400g at 4°C for 10 min. Cells were then fixed in 4% paraformaldehyde in PBS (PFA) for 10 min at RT, after which the fixation was stopped with 130 mM Glycine. 10% BSA was added in a 1:1 ratio and cells were pelleted by centrifugation (10 min, 400g, 4°C). Finally, cells were incubated in 30% sucrose/PBS solution at 4°C, scooped from tubes and embedded in freezing medium as described for the tissues.

### CUT&Tag

#### Sample preparation

Samples were prepared for CUT&Tag assay as described previously with several modifications^9,41^. For G4 mESCs, feeder cells were depleted from trypsinzed cell suspensions through their preferential adherence to sequential tissue culture dishes. Following centrifugation (5 min, 300g, RT), cells were resuspended in CUT&Tag wash buffer (20 mM HEPES pH 7.5 (Jena Bioscience, BU10675), 150 mM NaCl (Invitrogen, AM9760G), 0.5 mM spermidine (Sigma, S0266), 10 mM sodium butyrate (Sigma Aldrich, B5887), 1 mM PMSF (Thermo Scientific, 36978), Protease Inhibitor Cocktail (Roche, 04693132001)) and quantified using EVE automatic cell counter (NanoEntek).

For sampling rod cells, retinas from 2 month old NRL-GFP mice were dissociated using the Papain Dissociation System (Worthington, #130-094-802) as described above. Four retinas from two mice were used for one biological replicate. FACS sorting of single cells was performed on the BD FACS Aria II or Aria III Cell Sorter System with sample and plate cooling at 4°C. Forward-side scatter gating and GFP signal was used to sort intact Rod cells into 15 ml falcons pre-coated with BSA and containing 5 ml of wash buffer for approximately 1 h. After sorting, rod cells were counted using a haemocytometer.

#### Tagmentation

CUT&Tag was performed as described previously with minor modifications^9,41,42^. Briefly, 11 μl Concanavalin A beads (BioMag, 86057;) were equilibrated twice with 100 μl and then concentrated in 11 μl of binding buffer (20 mM HEPES pH 7.5, 10 mM KCL (Invitrogen, AM9640G), 1mM CaCl_2_ (Merck, 102382), 1mM MnCl_2_ (Thermo, J63150.AD)). 100,000 cells were then bound to the beads by incubating for 10 min at RT with rotation. Beads were subsequently separated on a magnet and resuspended in 100 μl CUT&Tag Antibody buffer (Wash buffer with 0.05% Digitonin (Milipore, 30410) and 2 mM EDTA (Invitrogen, 15575-038)). Subsequently, 1 μl of primary Ab (Supplementary Table 2) or IgG control (Cell Signaling, 68860 for mouse; Cell Signaling, 2729 for rabbit) was added and incubated on a rotator O/N (15 h) at 4°C. After magnetic separation, beads were resuspended in 100 μl of Dig-wash buffer containing 1 μl of matching secondary antibody (Abcam, ab46540 for mouse; Antibodies-online, ABIN101961 for rabbit) and incubated for 1 h at 4°C with rotation. Following three washes with CUT&Tag Dig-wash buffer (Wash buffer with 0.05% Digitonin), beads were resuspended in 100 μl CUT&Tag Dig-300 buffer (20 mM HEPES-KOH, pH 7.5, 300 mM NaCl, 0.5 mM Spermidine, 0.01% Digitonin, 10 mM Sodium butyrate, 1 mM PMSF) with a 1:250 dilution of homemade 3xFLAG-pA-Tn5 preloaded with Mosaic-end adapters. After incubation at 4°C for 1 h with rotation, beads were washed four times with Dig-300 buffer and resuspended in 50 μl Tagmentation buffer (Dig-300 buffer 10 Mm MgCl2 (Invitrogen, AM9530G)). Homemade spike-in-AmpR was added to the tagmentation buffer at a concentration of 33.6 fM^42^. Tagmentation was performed for 1 h at 37°C and subsequently stopped by adding 2.25 μl EDTA, 2.75 μl 10% SDS (Invitrogen, 15553-035) and 0.5 μl Proteinase K (Invitrogen, 25530049). DNA fragments were solubilized for 16 h at 55°C followed by 30 min at 70°C to inactivate residual Proteinase K. DNA fragments were finally purified and eluted with 25 μl of elution buffer (Zymo, D5205).

#### Library preparation and sequencing

NGS libraries were generated by PCR amplifying the CUT&Tag DNA fragments with barcoded i5 and i7 primers^43^ as described previously^41^. Following PCR, 1x volume of Ampure XP beads (Beckman Coulter) were mixed with the NGS libraries and incubated at RT for 10 min. After magnetic separation, beads were washed three times on the magnet with 80% ethanol, and the libraries were eluted with Tris-HCL, pH 8.0 (Roth, 9090.3). The quality of the purified NGS libraries was assessed with the D5000 tapestation (Agilent Technologies). The final libraries were sequenced on the Illumina NovaSeq X Plus platform with PE100 read length, yielding between 5 and 16 million fragments per sample.

#### NGS data processing

Sequencing data was processed as previously described with several modifications^42,44^. Briefly, sequencing adapters were trimmed using Trim Galore (v0.6.4) with the options --paired -- nextera^45^. The trimmed reads were then aligned to the mouse genome (mm39) using Bowtie2 (v2.3.5.1)^46^ with --local --very-sensitive-local --no-mixed --no-discordant --phred33 -I 10 -X 2000. Aligned reads were filtered for mapped, paired-end reads (MAPQ ≥ 20) using samtools (v1.10) with view -f 2 -q 20 and then subsequently sorted. ENCODE blacklisted regions^47^ were removed by bedtools (v2.29.2)^48^ with intersect -v. For visualization, normalized coverage tracks (bigWig) were generated with deepTools2 (v3.4.1)^49^ using a bin size of 10 bp, read extension (-e), and CPM normalization, ignoring chrX and chrY. For merged tracks FASTQ files were concatenated from biological and technical replicates after visual inspection and statistical tests confirmed they were sufficiently similar. Specifically, deepTools2 multiBigwigSummary and plotCorrelation functions were used for 500bp window and pearson correlation (v3.4.1)^49^.

To visualize enrichment relative to the IgG control, log_2_ ratio tracks were generated using deepTools bigwigCompare (v3.4.1)^49^. Analogous 20kb binned subtraction bedGraphs were then used for LAD calling as previously described^50^ with minor changes in parameters. Specifically, matrices were binarized with signal above a threshold of 0.1 considered positive (LAD) and below as negative (non-LAD). Adjacent LAD segments separated by gaps ≤50 kb were merged and, subsequently, regions smaller than 200 kb were removed. Finally, this LAD calling was used to calculate LAD signal enrichment of other samples in the set using plotEnrichment from deepTools2 (v3.4.1)^49^.

Processed tracks and LAD calling are available for visualisation at https://genome-euro.ucsc.edu/s/Chudzik/AbTrapping.

### Immunostaining

#### Cultured and isolated cells

The primary and secondary Abs used in this study are presented in Supplementary Table 2 and 3 respectively. Cells growing on coverslips were rinsed with PBS pre-warmed to 37°C and fixed with freshly made 4% formaldehyde, pH 7.0 for 10 min at RT. Cells were washed with PBS+0.01% Tween20, 3 × 10 min at RT and permeabilized with 0.5% Triton X-100 in PBS for 10 min at RT. After washing with PBS for 10 min, coverslips were stored in PBS at +4°C until use for up to one week. Primary and secondary Abs were diluted in blocking solution consisting of 4% BSA in PBS+0.01% Tween20. Incubation with Abs was performed in dark chambers by placing coverslips up-side-down on drops of Ab solution loaded on Parafilm. If not otherwise indicated, incubations with primary and secondary Abs were for 1 h at RT (23-25°C). For DNA counterstain, DAPI or Hoechst 33342 was added to secondary Ab to final concentration of 2 or 1 µg/ml. Washings between and after Ab incubations were performed with PBS+0.01% Tween20 pre-warmed to 37°C, 3 × 10 min each at 37°C. Coverslips were mounted on microscopic slides with antifade medium Vectorshield (VectorLabs) where excess antifade was removed with soft tissue and coverslip edges were sealed with colorless nailpolish. For epitope reduction, gentle protein digestion with Trypsin at 37°C for 10 min was applied with a concentration of 6.25 µg/ml (for H3K9me2) or 12.5 µg/ml (for B23).

The HA-PRRX1 fusion protein was detected with rabbit-anti-HA primary antibodies (Cell Signaling, #3724) diluted 1:100 as described before^37^.

#### Immunofluorescence analogous to the CUT&Tag

100,000 single cells resuspended in 1 ml CUT&Tag wash buffer were attached to poly-lysine coated coverslips for 60 min at 4°C. Subsequently, non-attached cells were removed by two 5 minute incubations at 4°C with the wash buffer. Incubation with Abs was performed in dark chambers by placing coverslips up-side-down on drops of Abs loaded on Parafilm. Primary antibodies were used in the same quantities, buffers, and incubation times as described for CUT&Tag (see above). Donkey-anti-Rabbit Alexa555 (Invitrogen, A31572) and donkey anti-Mouse Alexa647 (Invitrogen, A-31571) were used as secondary Abs at 1:500 dilution. DAPI was added to the secondary Ab to a final concentration of 2 µg/ml. After both primary and secondary antibody incubations, coverslips were washed with a CUT&Tag Dig-Wash buffer at 4°C 2 × 10 min each. After final wash, cells were postfixed with 4% PFA, PBS+0.01% Tween20, 2 × 10 min at RT, and mounted on microscopic slides, as described above for standard immunofluorescence.

#### Immunostaining of cryosections

Before staining, cryosections were dried out for 20 min at RT to fix sections on microscopic slides. For immunostaining of histone modifications, antigen retrieval was performed, which included re-hydration of sections in Na-citrate buffer for 5 min followed by heating up to 80°C in the same buffer for 15-30 min in a water bath. Sections were additionally permeabilized in PBS/0.5% Triton X100 for 1 h. Primary and secondary Abs were diluted in blocking solution consisting of PBS, 2% BSA, 0.1% saponin, 0.1% Triton X-100. Incubation with Abs was performed in dark wet containers under home-made glass chambers for 10-12 h at RT^51^. For nuclear counterstain, DAPI was added to secondary Ab to a final concentration of 2 µg/ml. Washings between and after Abs were performed with pre-warmed to 37°C PBS/0.01% Triton X-100, 3 × 30 min each at 37°C. Sections were mounted under coverslips with antifade and sealed as described above for cells.

### Microscopy and image analysis

#### Confocal microscopy

Confocal image stacks were acquired with a TCS SP5 confocal microscope (Leica) using a Plan Apo 63/1.4 NA oil immersion objective and the Leica Application Suite Advanced Fluorescence Software (Leica). Z step size was adjusted to an axial chromatic shift and typically was either 200 nm or 300 nm. XY pixel size varied from 20 to 50 nm. Axial chromatic shift correction, as well as building single grey-scale stacks, RGB-stacks, montages and maximum intensity projections were performed using the ImageJ plugin StackGroom^52^ available upon request. For Fig.3a, Fig.4a, Extended Data Fig.1a, and Extended Data Fig.2a, confocal image stacks were acquired with a A1 confocal microscope (Nikon) using Plan Apo lambda 100/1.45 NA oil immersion objective lens and the NIS-elements AR version 5.21.00 software (Nikon).

#### Time-lapse microscopy of immunostaining

MEFs grown on a 24-well glass-bottom plates (AGC Techno Glass) were fixed, permeabilized, and blocked with PBS containing 4% BSA and 0.01% Tween 20. Plates were then set on an inverted fluorescence microscope (Nikon, Ti-E;) equipped with a 100x Plan Apo VC (NA 1.4) oil-immersion objective lens (Nikon), a spinning disc confocal unit (Yokogawa, CSU-W1), a laser unit (Chroma Technologies Japan, LDI-7 Laser Diode Illuminator), and an EM-CCD (iXon+; Andor; gain multiplier 300) operated by NIS-elements software ver. 5.11.03 (Nikon). Immediately after antibody mixture was added to a well, time-lapse recording at room temperature was started using 405-, 470-, and 555-nm laser lines with a 405/470/555/640NIR dichroic mirror, and 520/60, 600/50, and 690/50 emission filters. For double staining using CMA317 and AM39239, 100 μl of a mixture containing CMA317 (20 μg/ml), AM39239 (1:100 dilution), Alexa Fluor 488-conjugated goat anti-mouse Fcγ (Jackson ImmunoResearch; PRID, AB_2338458, 20 μg/ml), Cy3-conjugated goat anti-rabbit IgG (H+L) (Jackson ImmunoResearch, PRID, AB_2337919; 20 μg/ml), and Hoechst 33342 (Nacalai Tesque, 4 μg/ml) was preincubated at room temperature for 2 h before adding to a well filled with 100 μL of PBS containing 4% BSA and 0.01% Tween 20. Fluorescence images were acquired every 5 min for 16.5 h.

#### Image quantification

Images were processed using a custom Fiji macro to automate nuclear detection and spatial signal intensity analysis on the mid optical section. For better nuclear segmentation, two or three channels were summarized. Background subtraction, thresholding and particle analysis were applied to detect nuclei and create regions of interest (ROIs). For each detected nucleus, concentric ring-shaped ROIs were generated by iteratively shrinking the initial ROI by 250 nm. Mean signal intensities were measured for all ring-shaped ROIs and visualized using Seaborn. The resulting plots display the mean intensity values for all measured samples per staining, normalized to the brightest signal in each channel. The y-axes represent the normalized mean signal intensity, error bars indicate the standard deviation across samples. The number of analyzed nuclei varied between 20 and 50 per staining.

#### Measuring relative H3K9me2-binding affinities in different antibodies

The relative affinities of two mouse monoclonal Abs, CMA317 and ab1220, were measured using ELSA. Briefly, a 96-well plate was coated with 1 μg/mL synthetic H3K9me2 peptide^53^ overnight, washed and blocked with PBS containing 0.1% Tween 20 and 1% BSA (PBST-BSA) for 1 h. After 3x washing with PBS containing 0.1% Tween 20, wells were incubated with a serial dilution series of antibodies (one-third dilution series from 0.1 μg/mL) in duplicate in PBST-BSA for 1 h, Cells were subsequently washed 3x in PBST, incubated with peroxidase-conjugated AffiniPure™ goat anti-mouse IgG (1:10,000; Jackson ImmunoResaerch; 115-035-003; RRID: AB_10015289) for 1h. After 3x washing with PBST, colorimetric assay was performed using 0.26 mg/mL o-phenylenediamine·2HCl in 0.1 M citrate buffer (pH 5.0) containing 0.01% hydrogen peroxidate by measuring absorbance at 450 nm.

To compare the binding affinity of AM39239 to CMA317, the concentration of the bound fraction to H3K9me2 was first estimated. This is because unpurified rabbit polyclonal antibodies (AM39239) contain bulk IgG that do not bind to the target H3K9me2, unlike mouse monoclonal antibodies. Briefly, Dynabeads™ MyOne™ Streptavidin T1 (ThermoFisher Scientific, 100 μl) wereincubated with 1 μg of biotinylated histone H3K9 or histone H3K9me2 peptide (Active Motif) for 1 h at room temperature. The beads were washed with PBST and mixed with rabbit polyclonal antibody AM39239 (12.5 μL beads per 1 μl antibody) for 2 h at room temperature. The supernatant was collected and the beads were washed three times with PBST and eluted using 1x SDS sample buffer without DTT (50 mM Tris-HCl [pH 6.8], 2% SDS, 10% glycerol, 0.01% bromophenol blue). Input and supernatant samples (0.05 μl equivalent) and 10x concentrated bead eluate were separated on a 7.5% polyacrylamide gel (SuperSep™ Ace, 17 well pre-cast; Fujifilm Wako Chemicals) and stained with Bullet CBB Stain One (Nacalai Tesque). Relative band intensities were measured using ImageJ (https://imagej.net/ij/). The absence of H3K9me2-biding fraction from the supernatant was confirmed by immunofluorescence. We then compared the binding efficiency of rabbit and mouse antibodies to H3K9me2 beads. 0.5 μl of the rabbit polyclonal antibody (AM39239; 0.125 μg of H3K9me2-specific IgG fraction) and the mouse antibody (CMA317; 0, 0.625, 2.5, and 10 μg) were mixed and incubated with Dynabeads™ MyOne™ Streptavidin T1 beads (1.56 μl equivalent; premixed with 0.0156 μg of biotinylated histone H3K9me2 peptide) for 2 h at room temperature with rotation. After the supernatant was collected, beads were washed twice with PBST. The input, supernatant, and bead fractions were denatured by heating at 95°C for 5 min in 1x SDS sample buffer without DTT, before separating on a 7.5% polyacrylamide gel. H3K9me2-binding activity in supernatants was assessed by immunofluorescence.

#### Computational modeling of antibody staining

We modeled antibody staining as a reaction-diffusion system of antibodies (A) and antigens (B) inside the nucleus which can be described by the following partial differential equations (PDEs),

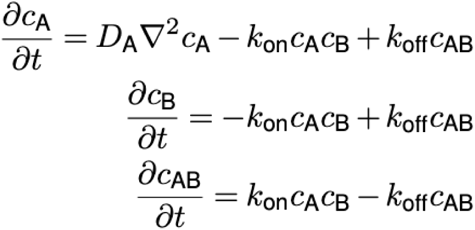

where c_A_(r, t), c_B_(r, t), and c_AB_(r, t) are the spatiotemporal concentration fields of antibody, antigen and antibody-antigen complex respectively. D_A_ is the diffusion constant of antibodies. The diffusion coefficient of the antigen (D_B_) and the antibody-antigen (D_AB_) complex were set to be zero, as the antigen considered is on histones which are bound to cross-linked chromatin. k_on_ and k_off_ are rate constants for the antibody binding and unbinding. For initial conditions, antigen was uniformly distributed in a sphere of radius R_0_=5 um at concentration c^0^_B_; antibodies were distributed outside of this sphere in a concentric shell of R_1_=75 um at concentration c^0^ . To mitigate the effect of stiffness for numerically solving the coupled equations, we smoothed the initial concentrations c_A_ (r, t=0), c_B_(r, t=0) by a hyperbolic tangent function and scaling factor ɛ was set to be 0.2:

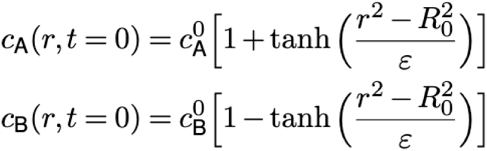

We discretized the problem in spherical coordinates, assuming spherical symmetry of the nucleus, which reduces the three-dimensional system to a one-dimensional problem, using a spatial grid spacing of 0.01 μm. To address the challenges of solving stiff equations when K_d_ (i.e., k_off_ / k_on_) is small, we utilized the Julia DifferentialEquations package, using the Rodas5P solver. This solver is a fifth-order A-stable Rosenbrock method, specifically designed for stiff problems, and is complemented by a stiff-aware fourth-order interpolant for improved accuracy and stability^54,55^.

## Data availability

All data reported in this paper are available upon request. Sequencing data generated in this study are available at the NCBI Gene Expression Omnibus, GEO: GSE293376. Processed CUT&Tag tracks are also available at https://genome-euro.ucsc.edu/s/Chudzik/AbTrapping for visual exploration in UCSC.

## Code availability

Code for computational modeling has been deposited at https://github.com/Fudenberg-Research-Group/Ab-trapping.

## Acknowledgements

We thank Corinna Schwichtenberg (MPIMG Animal house) for mouse line maintenance; former and current Schulz lab members (Alexandra Martitz, Till Schwämmle, Ilona Dunkel and Rutger Gjaltema) for kindly sharing an optimized CUT&Tag protocol; Marius Ader (TU Dresden) for kindly sharing NRL-GFP mouse strain; Bulut-Karslioglu lab for kindly sharing pA-Tn5; the MPIMG Transgenic unit for IVF; MPIMG FACS facilities for sorting; MDC/BIH Genomics Platform for sequencing. We are grateful to Evgenya Popova (Penn State Neuroscience Institute) and Sergei Grigoryev (Penn State College of Medicine) for fruitful discussions. COS-7 cells preparations with fusion protein PRRX1-HA were kindly donated by Rebecca S. Tooze (University of Oxford). X.Y. and G.F. thank members of the Fudenberg group for feedback on simulations. Finally, we thank Peter Becker, Silke Lochs, Philipp Kurbel and Robin van der Weide for their critical reading of the manuscript. K.C. was supported by the Deutsche Forschungsgemeinschaft (DFG) International Research Training Group (IRTG2403). Y.S. was supported by JST CREST (JPMJCR20S6). X.Y. and G.F. were supported by NIGMS (R35GM143116) to G.F. M.I.R. was supported by the EMBO (ALTF1554-2016), Wellcome Trust (206475/Z/17/Z) and the DFG grants (project # 556274722 and FOR 2841, project # 400728090). S.U. and I.S. were supported by the DFG grants (SP2202/SO1054/2, project # 422388934 and SFB1064, project # 213249687).

## Contributions

I.S., M.I.R., H.K, and K.C. conceived and designed the study. K.C. performed CUT&Tag experiments, including immunostaining of isolated cells, and data processing. X.Y. and G.F. developed computational models. W.T. performed antibody affinity measurements. S.U. performed immunostainings and image quantification. Y.S. and H.K. performed immunostaining and image acquisition in TT2 mESCs and 3KO/5KO MEFs, as well as time-lapse microscopy. L.S. performed image acquisition of COS-7 cells. I.S. performed immunostaining and image acquisition in both cultured cells and tissue sections. I.S., M.I.R., and K.C. wrote the manuscript with input from all authors.

## Corresponding author

Correspondence to Geoffrey Fudenberg, Hiroshi Kimura, Michael I. Robson or Irina Solovei.

## Ethics declarations

The authors declare no competing interests.

